# The exoproteome and surfaceome of toxigenic *Corynebacterium diphtheriae* 1737 and its response to iron-restriction and growth on human hemoglobin

**DOI:** 10.1101/2024.05.19.594877

**Authors:** Andrew K. Goring, Scott T. Hale, Poojita Dasika, Yu Chen, Robert T. Clubb, Joseph A. Loo

**Author notes:** To whom correspondence should be addressed: Robert T. Clubb, Department of Chemistry and Biochemistry, University of California, Los Angeles, 611 Charles Young Drive East, Los Angeles, CA 90095, USA, Joseph A. Loo, Department of Chemistry and Biochemistry, University of California, Los Angeles, 611 Charles Young Drive East, Los Angeles, CA 90095, USA.

## Abstract

Toxin-producing *Corynebacterium diphtheriae* strains are the etiological agent of the severe upper respiratory disease, diphtheria. A global phylogenetic analysis revealed that biotype gravis is particularly lethal as it produces diphtheria toxin and a range of other virulence factors, particularly when it encounters low levels of iron at sites of infection. To gain insight into how it colonizes its host we have identified iron-dependent changes in the exoproteome and surfaceome of *C. diphtheriae* strain 1737 using a combination of whole-cell fractionation, intact cell surface proteolysis, and quantitative proteomics. In total, we identified 1,425 of the predicted 2,265 (63%) proteins encoded by its reference genome. For each protein we quantified its degree of secretion and surface-exposure, revealing that exoproteases and hydrolases predominate in the exoproteome, while the surfaceome is enriched with adhesins, particularly DIP2093. Our analysis provides insight into how components in the heme-acquisition system are positioned, showing pronounced surface-exposure of the strain-specific ChtA/ChtC paralogues and high secretion of the species-conserved heme-binding HtaA protein suggesting it functions as a hemophore. Profiling the response of the exoproteome and surfaceome after microbial exposure to human hemoglobin and iron limitation reveals potential virulence factors that may be expressed at sites of infection. Data are available via ProteomeXchange with identifier PXD051674.

## INTRODUCTION

Diphtheria is a severe upper respiratory disease that can cause fatal infections.^1–4^ There are four biotypes of *C. diphtheriae*: gravis, mitis, intermedius and belfanti.^2^ Both toxigenic (encoding diphtheria toxin, “DT”) and non-toxigenic *Corynebacterium* subspecies are capable of colonizing the upper respiratory tract in humans.^1,5^ However, a global phylogenetic analysis suggests biotype gravis predominated in the Eastern European 1990s outbreak and was particularly lethal because it had re-acquired the DT gene (*tox*) from a β corynephage.^6^ Genomic sequencing of the related NCTC13129 strain subsequently revealed the presence of pathogenicity islands that contain a number of genes that encode for surface and secreted virulence factors, including pili,^7,8^ heme-acquisition machinery,^7,9,10^ and antimicrobial-resistance genes.^3,6^ Many non-pathogenic *Corynebacterium* subsp. are also part of the nasal commensal microbiota and are known to attenuate the virulence of more lethal opportunistic pathogens such as *Staphylococcus aureus*.^2^ Thus, defining the full complement of secreted (exoproteome) and surface (surfaceome) protein factors produced by *C. diphtheriae* that enables it to colonize its host promises to provide insight into how it both causes and prevents disease.

*C. diphtheriae* belongs to the Actinomycetota (or Actinobacteria) phylum, a group of high guanine-cytosine (G-C) gram-positive bacteria that contain other pathogenic species such as *Mycobacterium tuberculosis* and *Nocardia nova*.^11,12^ This large group of terrestrial and aquatic microbes are a unique hybrid of mono-and di-derm bacteria because their plasma membrane is surrounded by a thick peptidoglycan cell wall that is covalently modified by mycolic acid-linked secondary cell wall arabinogalactan polymers that form an outer myco-membrane (**Fig 1A**).^13^ This mycolyl-arabinogalactan-peptidoglycan (mAGP) complex prevents osmotic lysis and is highly impermeable, limiting the utility of some antibiotics to treat infections caused by Actinobacteria.^14^ For example, penetration of β-lactam drugs through the mycobacterial cell wall occurs 100-fold slower than through *Escherichia coli*’s cell wall.^15,16^

**Figure 1:**
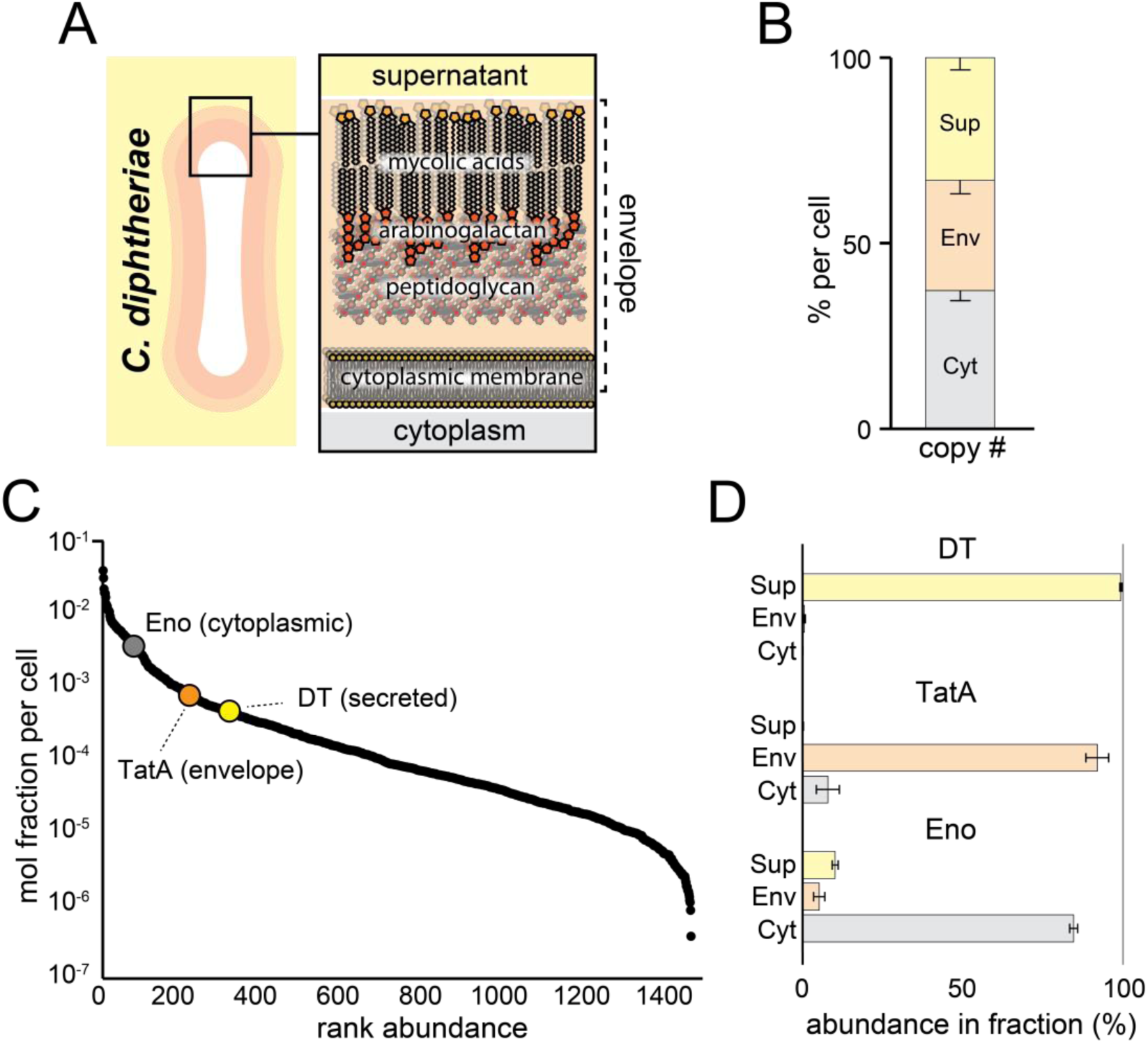
Cell fractionation of *C. diphtheria* grown in low iron media. (A) Diagram depicting the surface structure of *C. diphtheriae* and cellular fractions studied by label-free quantitative proteomics (supernatant, envelope, and cytoplasmic fractions). The envelope fraction presumably contains proteins within the plasma membrane, peptidoglycan, arabinogalactan, and mycolic acid layers. (B) Total abundance (in molar units) of all proteins found in each fraction expressed as a percentage of the total abundance of proteins found per culture, calculated using iBAQ values from MaxQuant^60^ (see methods). (C) Rank abundance plot for each protein. The abundance for each protein is its average total mole fraction per cell, obtained by summing its content in the cytoplasmic, supernatant and envelope fractions. The identities of three representative proteins for the secreted, envelope, and cytoplasmic fractions are labeled: diphtheria toxin (DT, yellow), twin arginine transporter protein A (TatA, orange), and enolase (Eno, gray). A criterion for a protein to be included in the analysis was that it was identified in each of the three biological replicates. (D) Localization profiles for the three representative proteins indicated in panel (C). The profile illustrates the relative % partitioning of each protein into the cytoplasmic (Cyt), envelope (Env), and secreted (Sup) fractions. Error bars are the standard deviation obtained from three biological replicates.

To effectively interact with their environment, the exoproteome and surfaceome in bacterial pathogens contain a range of proteins that promote among other functions, cell envelope maintenance, nutrient acquisition, and microbial adhesion to abiotic surfaces, other bacteria and host tissues.^17^ The first *C. diphtheriae* proteomic study was performed by Hansmeier and colleagues in 2006 who catalogued 85 proteins from supernatant and surface extracts produced by the non-toxigenic laboratory strain, *C7_s_*(-)*^tox-^*.^18^ A highly abundant papain-like cysteine peptidase (RipA, also known as RpfI) was identified that was later shown by Burkovski and colleagues to be critical for epithelial cell adhesion and internalization.^19^ In subsequent proteomic studies, highly abundant trypsin-like exoproteases were observed in a non-toxigenic strain (str. ISS3319)^20^ and other *Corynebacterium* subspecies.^21–23^ These enzymes may degrade host tissue or promote the maturation of extracellular virulence factors.^24,25^ Others have shown that surface adhesins such as pili play a role in colonization and virulence,^2^ as genetic elimination of the SpaA-pilus in pathogenic strain NCTC13129 decreased its lethality in a mouse model for infection.^26^ Interestingly, a quantitative proteomic analysis focusing on ISS13129 whole-cell lysates showed pathogenicity seems to be linked to nutrient poor growth conditions.^20^ Furthermore, due to metal-ion sequestration experienced during infection,^27^ surface-displayed and secreted virulence factors are employed by pathogenic bacteria as part of a systems-level response termed metallostasis, which aid not only in metal scavenging, efflux, and intracellular sequestration, but also in ribosome remodeling and metabolic reprograming.^28^ Work by Schmitt and colleagues has identified several surface components in the system (HtaA, HtaB, ChtA, ChtB, ChtC, and HbpA),^29–31^ which facilitate growth on human hemoglobin (Hb) and its complex with haptoglobin^10^ to overcome host innate nutritional immunity mechanisms.^32–35^ A proteomic analysis on the vaccine – essentially crude purifications of inactivated DT isolated from the supernatants of high *tox*-producing strains (e.g. str. PW8) cultured under low-iron conditions – have shown that iron limitation triggers the expression of DT^32,36–38^ and several other immunogenic proteins.^39^ However, despite its clinical relevance and use as a model bacterium for research purposes, no proteomic study reported to date has rigorously defined the exoproteome and surfaceome in any *tox*-producing *C. diphtheriae* strain or defined how these proteomes change in response to iron limitation.

In this study we determined how *C. diphtheriae* strain 1737 alters its exoproteome and sufaceome in response to iron limitation, a key microbial nutrient that is actively scavenged during infections and whose abundance is known to regulate the secretion of DT.^34^ Using a combination of cell fractionation and surface shaving methods, we determined the preferential localization and abundance of secreted and surface-exposed proteins. We found highly abundant exoproteases (DIP2069, RipA) and resuscitation-promoting factors (RpfA, RpfB) are preferentially secreted into the exoproteome, while the surfaceome is enriched with primarily adhesins (DIP2093) and corynemycolic acid transferases that that maintain the structure of the outer myco-membrane (cMytA, cMytB, cMytC). Our results also provide a spatial view of how *C. diphtheriae*’s heme-acquisition system components are distributed between the microbial surface and exoproteome, suggesting a possible hemophore function by HtaA which is predominantly localized to the exoproteome. Finally, we show that iron-restriction causes significant changes in a subset of exoenzymes, cell envelope maintenance components, and uncharacterized proteins in addition to DT, microbial siderophores-uptake components, hemophores. Interestingly, the source of iron, either heme from Hb or free iron from FeCl_3_, also affects the abundance of key hemophores that are transcriptionally regulated by the HrrAS and ChrSA two-component systems.^40,41^ Together, our results provide a comprehensive view of the exoproteome and surfaceome in *C. diphtheriae*, and they reveal how it changes when the microbe encounters iron-restricted conditions that are present at sites of infection.

## METHODS

### Strains and growth conditions

*C. diphtheriae* strain 1737 was provided by Dr. Michael Schmitt (Laboratory of Respiratory and Special Pathogens at the Federal Drug Administration). A single colony of *C. diphtheriae* from a heart infusion (HI) agar plate was inoculated into HI broth supplemented with 0.2% Tween-80 (HIBTW) and grown overnight at 37°C with agitation (200 rpm). The following day this culture was diluted 1:2 with HIBTW and grown for 1 hour before being washed and resuspended in iron-free chemically defined media, mPGT.^42^ This was used to inoculate 10 mL cultures at a starting OD_600_ of 0.1 for all proteomics experiments. The four growth conditions used for proteomics experiments in this study were mPGT medium with 0.3 μM FeCl_3_ (“low-iron”, condition 1), 40 μM FeCl_3_ (“high iron”, condition 2a), 10 μM iron chelator EDDHA (ethylenediamine-N,N′-bis(2-hydroxyphenylacetic acid)) + 10 μM FeCl_3_ (“Fe-restricted + FeCl_3_”, condition 2b), or 10 μM EDDHA + 0.3 μM adult human methemoglobin purified from blood (“Fe-restricted + Hb”, condition 2c).

### Isolation of secreted, envelope, and cytoplasmic proteins

Log-phase low-iron cultures (condition 1) were harvested by centrifugation (6,000 rcf, 6 minutes); the supernatant was separated from the cell pellet and centrifuged again with protease inhibitor cocktail (Promega) (6,000 rcf, 60 minutes, 4 °C) and filtered with a 0.2 µm filter (Millipore Sigma) to ensure it was cell-free. The extracellular proteins in the filtrate were then concentrated in a 10kDa MWCO Amicon (Millipore Sigma) centrifugal filter. A subset of the resulting sample was quantified using the bicinchoninic acid (BCA) assay (Pierce) following a 4-hour dialysis into lysis buffer (PBS, pH 7.3) to eliminate background effects from the media. The cell envelope and cytoplasmic fractions were prepared from the cell pellet. Pelleted cells were washed three times with wash buffer (0.2% sodium azide + PBS, pH 7.3) and an aliquot of 2 OD units (OD600*mL) were saved for surface shaving experiments (described below). The remaining cells were resuspended in 1 mL lysis buffer + protease inhibitor cocktail (Promega) and sonicated using a microtip sonicator for 8 minutes total (8 rounds: 60s on, 30s off) in a microcentrifuge tube soaked in a circulating ice water bath. Following lysis, cell debris and unlysed cells were removed from the whole cell lysate by centrifugation at 3,000 g, 4°C, for 15 minutes; the OD600 of the pellet resuspended in 1 mL lysis buffer was used to calculate lysis efficiency (described below). The whole cell lysate (supernatant) was centrifuged again at 22,000 g, 4°C, for 1 h to yield the soluble cytoplasmic fraction and cell envelope pellet. The cell envelope fraction was resuspended with lysis buffer and washed three more times. Supernatant, cytoplasmic, and cell envelope fractions were prepared for LC-MS/MS using enhanced filter-aided sample preparation (eFASP).^43^ Briefly, 50-100 ug from each fraction were buffer-exchanged into exchange buffer (8M urea, 0.1% deoxycholic acid, 0.1% β-octyl-glucoside, 100 mM ammonium bicarbonate, pH 7.8) using a 10 kDa MWCO amicon filter, Cys-alkylated, and digested overnight with a 1:50 ratio of protein:trypsin (Promega, sequencing grade). Detergents were removed through organic-aqueous phase extraction with ethyl acetate, and samples were dried using a speed vac, before performing a stage-tip C18 desalting procedure.^44^

### Intact cell surface proteolysis

Surface shaving was performed on intact cells as described in the literature^45^ with minor modifications as described here. Log-phase low-iron cell pellets were washed three times with wash buffer and 2 OD units were resuspended in 1 mL PBS (pH 7.3).

Each biological replicate was split into two 500 uL aliquots and incubated with either 3 µg of trypsin (Promega, sequencing grade) (“surface digest”) or protease-free buffer (tris HCl, pH 7.8) (“control”) for 15 minutes at 37 °C with gentle shaking. Each incubation was pelleted (13,000 rcf, 10 minutes, 4 °C) and the supernatant was collected and filtered using a 0.2 µm filter (Millipore Sigma). This aliquot was quantified using BCA assay and subject to disulfide reduction, Cys-alkylation with iodoacetamide, and an overnight digest with trypsin to complete digestion of the released peptides at a 1:50 ratio at 37°C and desalted using a stage-tip C18.^44^

### LC-MS/MS

Each sample was injected to an Ultimate 3000 nano LC, which was equipped with a 75 μm x 2 cm trap column packed with C18 3 μm bulk resins (Acclaim PepMap 100, Thermo Scientific) and a 75 μm x 15 cm analytical column with C18 2 μm resins (Acclaim PepMap RSLC, Thermo Scientific). The nanoLC gradient was 3−35% solvent B (A = H_2_O with 0.1% formic acid; B = acetonitrile with 0.1% formic acid) over 40 min and from 35% to 85% solvent B in 5 min at flow rate 300 nL/min. The nanoLC was coupled to an Exploris 480 orbitrap mass spectrometer (Thermo Fisher Scientific, San Jose, CA). Full spectra (*m/z* 350 - 2000) were acquired in profile mode with resolution 120,000 at *m/z* 200 with an automatic gain control (AGC) target of 3 × 10^6^ every second. The most abundant ions were subjected to fragmentation by higher-energy collisional dissociation (HCD) with a normalized collisional energy of 30. MS/MS spectra were acquired in profile mode with resolution 15,000 at *m/z* 200 and AGC target of 1 × 10^6^. Charge states 1, 7, 8, and unassigned were excluded from tandem MS experiments. Dynamic exclusion was set at 45.0 s.

### Mass spectrometry data analysis

Raw MS data was searched against the *C. diphtheriae* database (Uniprot entry: UP000002198, accessed December 2023) using MaxQuant. The following parameters were set: precursor mass tolerance ± 20 ppm, fragment mass tolerance ± 0.02 Da for HCD, up to two missed trypsin cleavages, methionine oxidation as variable modification. The false discovery rate was 1.0% and at least 1 peptide was required for protein identification. Only proteins that were identified in at least three biological replicates of at least one fraction were included in our analysis. Only proteins with at least two peptides in either the surface digest or supernatant fractions (in all biological replicates) were used to evaluate preferential secretion or surface exposure.

### Protein quantification and construction of localization profiles

Protein samples were quantified using the BCA assay on the (1) cytoplasm, (2) envelope, (3) supernatant, and (4) surface digest. For fractions 1-3 above, total protein content per culture was calculated according to equation 1 below, using the concentration determined by the BCA assay and the volume of the soluble fraction for (1), resuspension of the membrane pellet for (2), and concentrated supernatant for (3). Lysis efficiency (equation 2) was determined based on the optical density at 600 nm for the sample before lysis and cell debris resuspended to an equal volume. All three biological replicates had consistent lysis efficiency of approximately 70% (or 0.7 in the equations below) which used for envelope and cytoplasmic fractions and 1.0 was used for the supernatant.

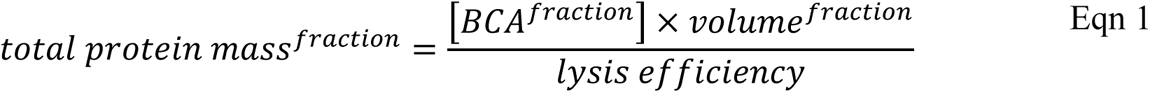

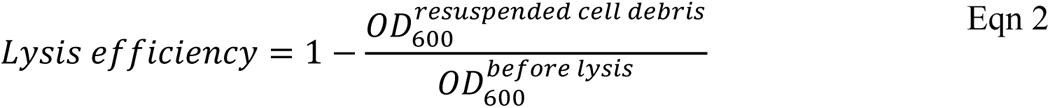

The percent copy number of each fraction per culture and construction of localization profiles was performed using a variation of the total protein approach (TPA).^46,47^ This relies on the intensity-based absolute quantification (iBAQ) values from MaxQuant^48^ and the total BCA-quantified protein abundance per culture and fraction. Since iBAQ values report on the molar abundance of a protein, the relative mol fraction of each protein in a sample (riBAQ) can be calculated by normalizing its iBAQ value to the sum of those from the sample, excluding contaminants (equation 3). We then determined the average molecular weight of each sample from the sum of each protein’s molecular weight multiplied by its riBAQ, which was used to obtain molar quantities of protein per sample according to equation 4. The percent copy number for each fraction (**Fig. 1B**) was calculated by dividing the molar abundance of the fraction by the sum of that from all fractions of that replicate. Finally, for each biological replicate, we determined the percent abundance for each protein that came from the cytoplasmic, envelope, and supernatant fractions (equation 5). The averages and standard deviations of the percent abundances across biological replicates give our final localization profiles.

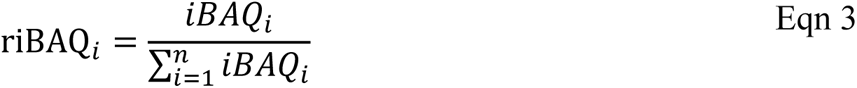

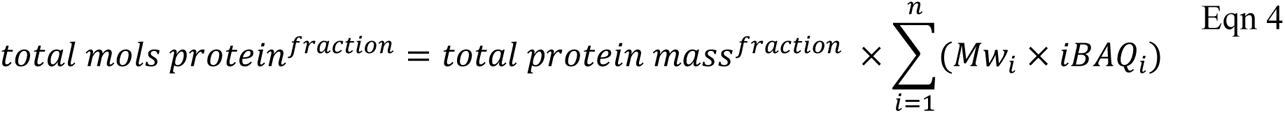

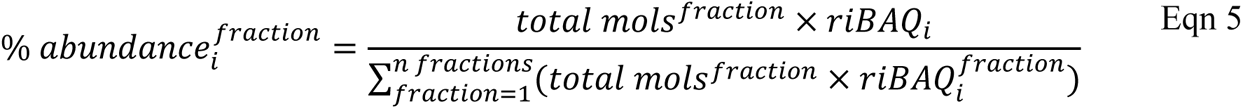

### Quantifying surface exposure

To determine the degree of surface exposure, we compared the protein intensities released from the surface digest to that of the whole-cell lysate (envelope + cytoplasmic fractions) after proper normalization by protein released from each fraction. Specifically, this involved first calculating the relative intensity of each protein in each fraction, which reports on the % of protein (by mass) in a sample. We used culture-normalized intensities rather than the iBAQ values used to generate localization profiles (**Fig. 1D**) because the calculation for iBAQ assumes proteins are intact, while the surface shaving approach is expected to release protein fragments that are exposed on the cell surface. The relative intensity of each protein was then scaled by the mass in grams of that sample for the envelope and surface digest fractions, respectively. The average ratio of the normalized surface digest intensities divided by whole-cell lysate intensities were used to rank surface exposure. The only protein whose average surface-exposure was greater than 1 was SpaB (DIP2011), which is highly abundant (30^th^ most abundant protein) from the surface digest fraction. While SpaB was identified from cellular fractions across all three biological replicates, only replicate three had at least two peptides identified in both the cytoplasmic and envelope fractions where its surface-exposure was 95%, suggesting this more accurate reading is a better representation of its true ratio. Thus, we assigned it an average value of 100% as the larger ratios are likely due to experimental error. Small ribosomal subunit protein bS16 (RpsP) was also highly abundant and not detected from the control incubation but is likely cytoplasmic as more than half of its abundance stemmed from the cytosolic fraction. A handful of ribosomal proteins that were discovered to have moderate and low preferential surface-exposure, but not in the predominantly surface-exposed category, were also removed if their localization profile had moderate or higher cytoplasmic localization or were identified from the control incubation.

### Bioinformatic analyses

We used 4 servers that look for sequence motifs either indicating secretion or presence outside of the cytoplasm. These are as follows: (1) the SignalP 6.0^49^ server to identify canonical signal sequences that indicate whether a protein is secreted through the Sec translocon (SP, signal sequences) or Twin-Arginine Translocation (TAT) pathways, and whether it also contains motifs associated with lipoproteins that are likely exported through these pathways, indicated by SPLIPO and TATLIPO, respectively; (2-3) PHOBIUS^50^ and DTMHMM^51^ servers that find transmembrane helices; (4) the CWpred^52^ server that looks for LXXTG-like sorting signals which indicate covalent attachment to the cell wall by sortase transpeptidases. The numbers of proteins encoding these sequences and their localization assignments as determined from our low-iron exoproteome and surfaceome analysis are summarized in supplemental **Figure S5**. Bioinformatic functional annotations such as domains and protein families were obtained for each protein from the most recent release of the eggnog 6.0 database^53^, Pfam,^54^ and InterproScan.^55^ Sequence homologues for *M. tuberculosis* strain H37Rv and their annotations were found by uploading the sequences of proteins identified in our study into String^56^ and exporting the closest H37Rv homologues.

## RESULTS

### Establishing growth conditions for proteomics studies

A total of four iron conditions were used in our proteomics experiments. Preliminary *C. diphtheriae* growth studies in iron-free chemically defined media (mPGT)^42^ revealed that a concentration of 0.3-40 µM FeCl_3_ in the culture media supports bacterial growth, above which growth is either partially or strongly reduced (**Fig. S1A**). Cells grown in low-iron media (0.3 µM FeCl_3_ in mPGT; “low-iron”, condition 1) were used for whole-cell fractionation and quantitative proteomic evaluation of the exoproteome and surfaceome. This condition has been used in the literature to study *C. diphtheriae* virulence-properties.^10,30^ Three more similar conditions (2a-2c, defined below) were used to study the response of the exoproteome and surfaceome to iron concentration and source. Condition 2a (“high-iron”; 40 µM FeCl_3_ in mPGT) contained the highest concentration of iron before growth was diminished growth, likely due to reactive oxygen species as seen in other bacteria.^57^ In contrast, the other two conditions utilized iron-restricted media that contained either FeCl_3_ (“Fe-restricted + FeCl_3_”, condition 2b) or human adult hemoglobin (Hb) purified from blood (“Fe-restricted + Hb”, condition 2c) as the sole source of iron. Each iron-restricted condition contains 10 µM iron-chelator (EDDHA), which completely abrogates growth unless the cells are supplemented with either 0.3 µM Hb (on a heme-iron basis) or 10 µM FeCl_3_ (**Fig. S1B**). We used iron-restricted conditions containing the EDDHA iron-chelator rather than simply adding Hb to deferrated growth media to more rigorously remove iron that can otherwise be leached from an array of sources (e.g. plastic and glassware). These more stringent iron-restricted conditions enable the effect of iron source to be accurately determined.

### Identification of secreted and cell envelope proteins in C. diphtheriae grown in low-iron media

Cells were grown in low-iron media (condition 1) to mid-log phase and then fractionated into cytoplasmic, cell envelope, and secreted protein fractions (**Fig. 1A**), which contain respectively 28.5±1.5%, 36.8±3.5%, and 34.7±3.8% of the total protein content by mass of the cell as determined by a BCA assay (**Fig. S2**). A combined total of 1,425 distinct proteins were identified from all three fractions, which represents 63% of the proteins encoded by the NCTC13129 reference genome (2,265 proteins; Uniprot proteome ID: UP000002198).^58^ A total of 1,100, 1,275, and 765 proteins were identified in the cytoplasmic, envelope, and secreted (supernatant) fractions, respectively. The resulting data was quantified using MaxQuant’s intensity-based absolute quantification (iBAQ) values,^48^ which report on protein abundance on a per-mole basis. Based on iBAQ values and amount of protein released from each fraction, we determined the relative copy number of proteins from each fraction (**Fig. 1B**) (see methods). The combined abundances show the mole fractions of identified proteins span just over 4 orders of magnitude (**Fig. 1C**). Two of the three most abundant proteins were ribosomal protein L7/L12 (RplL) and elongation factor Tu (Tuf), which is consistent with a prior proteomic analysis of whole cell lysate from non-toxigenic *C. diphtheriae* strain ISS3319.^5^ The second most abundant protein, a putative secreted protease (DIP2069), was predominantly found in the supernatant. The relative abundance of each protein in the different fractions was estimated by constructing its “localization profile” (see methods). The profile reveals a protein’s abundance in each fraction (cytoplasmic, envelope, or secreted), expressed as a percentage of its total abundance. Example protein localization profiles are presented in **Figure 1D**. Notably, the location profile for the diphtheria toxin (DT) indicates that it is predominantly secreted in low-iron conditions, in agreement with the literature.^59^ Expected localization profiles are also observed for other reference proteins, including TatA and enolase which are primarily located in the envelope and cytoplasmic fractions, respectively. The localization profiles for the most abundant proteins from each fraction are shown in **Fig. S3** and provided in **Table S1**. We implemented the localization profiles to quantify the degree of secretion for 436 of 765 proteins identified in the supernatant, for which at least two peptides were observed in each of the biological replicate supernatants. A protein’s localization preference for a particular fraction was defined as “predominant”, “moderate”, or “marginal” if >90%, >50-90%, or 30-50% of its total abundance was present in that fraction, respectively. Using this criterion, 32, 51, and 55 proteins had a predominant, moderate, and marginal preference for this fraction, respectively (**Fig. 2A**). The fourteen most abundant proteins in the exoproteome account for ∼40% of its total amount by mass and are illustrated in **Figure 2B**. Finally, there are 4 abundant proteins in the secreted fraction whose presence may occur as a result of cell lysis as they are highly abundant and predominantly found in the cytoplasm (**Figs. 2B, S2** RplL, ptsH, GroS, RplX). All proteins in the exoproteome, their abundances and preferential secretion are provided in **Table S2**.

**Figure 2:**
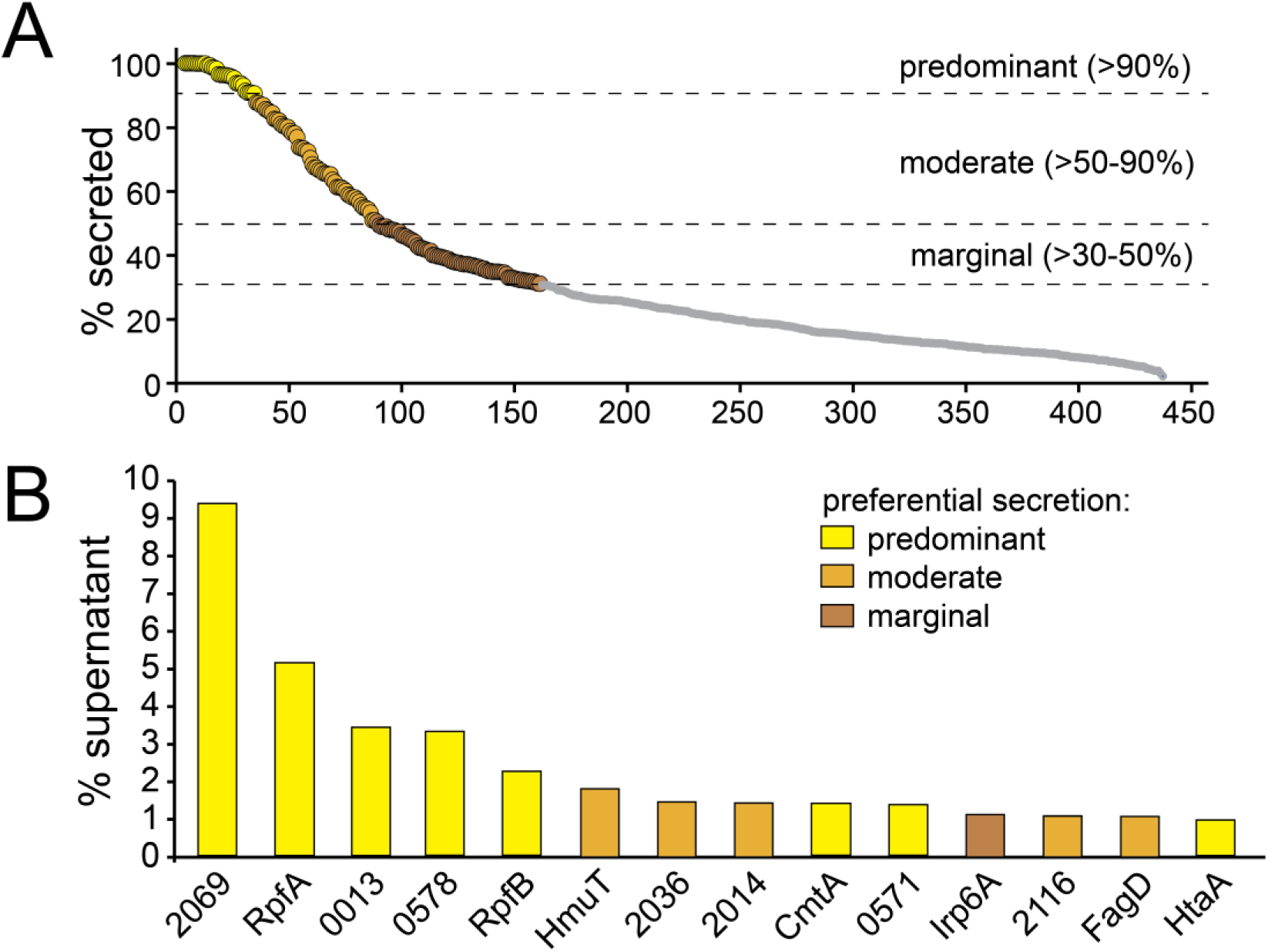
Highly abundant proteins in the supernatant, envelope and cytoplasmic fractions of cells grown in low iron media. (A) Plot of % secretion for all proteins quantified in the supernatant fraction, defined as the percent of the protein’s total culture-normalized abundance (from the supernatant, envelope, and cytoplasmic fractions) found in the supernatant. Secretion levels were defined as predominantly, moderately, and marginally secreted for proteins whose abundances are >90%, >50-90%, and >30-50% from the supernatant, respectively (dotted lines); those less than 30% secreted are indicated in grey and were not considered as part of the exoproteome. At least two peptides for a protein identified must have been present in the supernatant in all three biological replicates to be included in the analysis for quantification. Proteins in the exoproteome required a statistically higher fractional abundance in the supernatant relative to the cytoplasmic fraction (determined by t-test across all three biological replicates with a p-value <0.05) and at least 30% secreted; these proteins and their abundance from each fraction are listed in **Table S2**. (B) Bar graph showing the most abundant proteins from the supernatant with the y-axis showing the abundance of that protein relative to all proteins quantified in the supernatant, expressed as a percentage. Each protein is colored based on its preferential secretion, indicated on the graph.

### Identification of surface-exposed proteins in low-iron media

Surface-exposed proteins were identified by ‘surface shaving’ following a procedure in which cells grown to log-phase in low-iron media were harvested, washed and then subjected to limited trypsin digestion for label-free quantitative proteomics (**Fig. 3A**).^21,61,62^ A protease exposure time of 15 minutes was found to be optimal, as it yielded the largest differences in the amount of liberated peptides relative to an experimental control in which the same procedure was followed, but no protease was added (**Fig. S4**). Surface-exposure was calculated only for proteins that did not appear in the protease-free control and/or proteins that are not highly abundant in the cytoplasmic fraction (≤ 40% of its total abundance was in the cytoplasm) (see methods). To determine each protein’s degree of surface exposure, we calculated the ratio of its surface abundance in the shaving experiment relative to its summed abundance from the whole-cell lysate (envelope and cytoplasmic fractions, see methods). The total mass of protein released from the brief 15-minute proteolysis was 1.8 ± 0.4% of the total cellular protein content (envelope + cytoplasm), while the protease-free control incubation released 0.5 ± 0.3%. To reveal their degree of surface-exposure, proteins were ranked based on the ratio of their surface-proteolyzed abundance relative to their whole-cell abundance; a value of 1 corresponds to 100% of the protein being liberated from the brief 15-minute proteolysis (**Fig. 3B**). We defined “predominant”, “moderate”, and “marginal” levels of surface exposure for a protein if, respectively, >8.1%, >1.9-8.1%, and >0.2-1.9% of its total envelope + cytoplasmic intensity was released after surface proteolysis. A total of 25, 50, and 130 proteins were found to have predominant, moderate, and marginal levels of exposure, respectively. These surface proteins are listed in **Table S3** along with their fractional abundances used to generate localization profiles. The most abundant exposed proteins are putative adhesins (e.g. DIP2093, DIP2062), enzymes that maintain the outer membrane (corynemycolic acid transferases cMytA and cMytB), proteins involved in iron and heme import (e.g. Irp6A, ChtA, HbpA), and an uncharacterized protein, (DIP0659) (**Fig. 3C**). Notably, the putative adhesin DIP2093 is by far the most abundant protein on the cell surface (it is at least ∼4-times more abundant than any other protein). A handful of ribosomal proteins were also detected in the surface shaving experiments, but their presence is presumably an artifact as they are highly abundant and based on their localization profiles they primarily reside within the cytoplasm.

**Figure 3:**
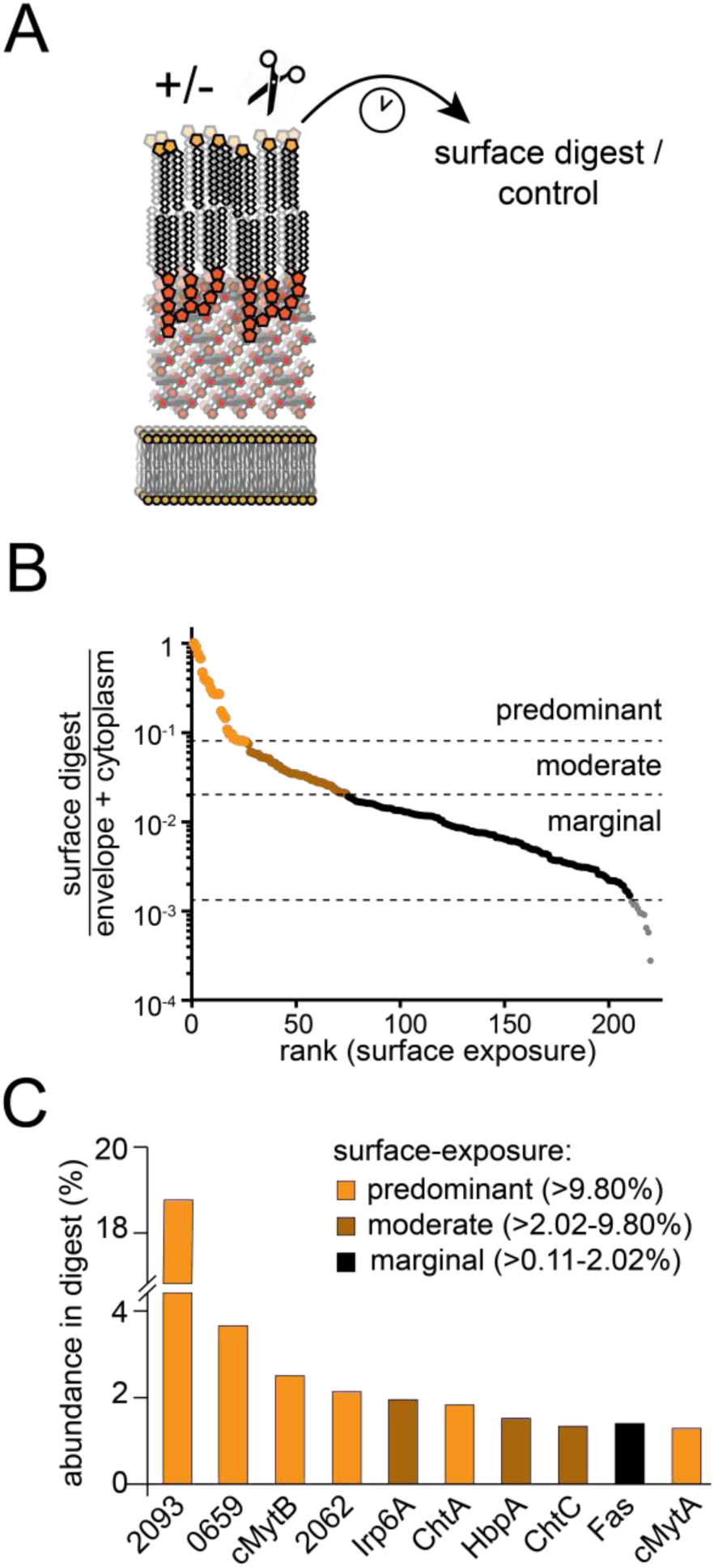
Identification of surface-exposed proteins. (A) Schematic depicting surface digest procedure using trypsin on intact cells. (B) Ranking of digest abundances over the cellular fractions (envelope & cytoplasmic) informs on relative surface exposure of proteins liberated from the surface. The degree of surface-exposure is assigned based on the percent of culture-normalized protein intensity released from the surface digest and defined as predominantly (>9.8%), moderately (>2.0-9.8%), and marginally (>0.1-2.0%) surface-exposed. In order for a protein to be assigned to the surfaceome, at least 0.11% of its culture abundance had to be released from the 15 minute surface proteolysis and proteins identified from the control incubation without a canonical secretion signal determined by SignalP6.0^49^ excluded; these are listed in **Table S2**. (C) The most highly abundant proteins in the surfaceome are labelled according to their preference for surface-exposure.

### Bioinformatics analysis of the surfaceome and exoproteome

We used bioinformatics approaches to predict the functions of proteins in the exoproteome and surfaceome. Proteins with predominant to moderate secretion or surface-exposure were analyzed for cells grown in low iron media (**Fig. 4A**). Specifically, we determine the abundances of “superfamily” and “family” type classifications from InterPro Scan^55^, which typically describe sequence, folds, or functions of a common evolutionary origin (**Fig. 4B**); similarly, for each protein we quantified the abundance of particular “domains” and “sequence repeats” to gain insight in their function (**Fig. 4C**). This analysis reveals that the exoproteome contains a large number of exoenzymes, with a majority originating from the trypsin-like superfamily of proteases (DIP2069, DIP0736, DIP0743, SprX, PepD proteins from the PA clan), lysozyme-like superfamily (RpfA, RpfA, Pbp1A, Pon1), α/β hydrolase fold superfamily (cMytA, cMytB, cMytC, ElrF, LpqC, DIP1119, DIP2339), and papain-like cysteine peptidases (RipA, RipC, RipC’ proteins from the CA clan) (**Fig. 4B**). The most abundant exoenzyme is a trypsin-like protease, DIP2069, which accounts for ∼10% of the secreted fraction (**Fig. 2B**). Highly abundant functional domains other than those associated with the aforementioned exoenzymes include the heme-binding conserved region (“CR”) domain (referred to as Htaa domains in the InterProScan analysis), and the ABC transporter periplasmic binding proteins involved in either heme-iron acquisition (HmuT) and iron-siderophore uptake (FrgD) (**Fig. 4C**). The surfaceome is dominated by many proteins with immunoglobulin-like folds (SpaB, DIP2093, DIP2062, DIP0357) and members of the α/β hydrolase fold superfamily (cMytA, cMytB, LipY, DIP2339). The most abundant protein is a large 101 kDa putative cell wall-anchored Sdr-family related adhesin, DIP2093 (it accounts for ∼18% of the total surface proteolyzed peptide intensity) (**Fig. 3C**). Additionally, proteins containing LGFP tandem repeats (cMytA, Csp) were also found to be prevalent in both the surface and secreted fractions (**Fig. 4C** and **Table S2**, **S3**).

**Figure 4:**
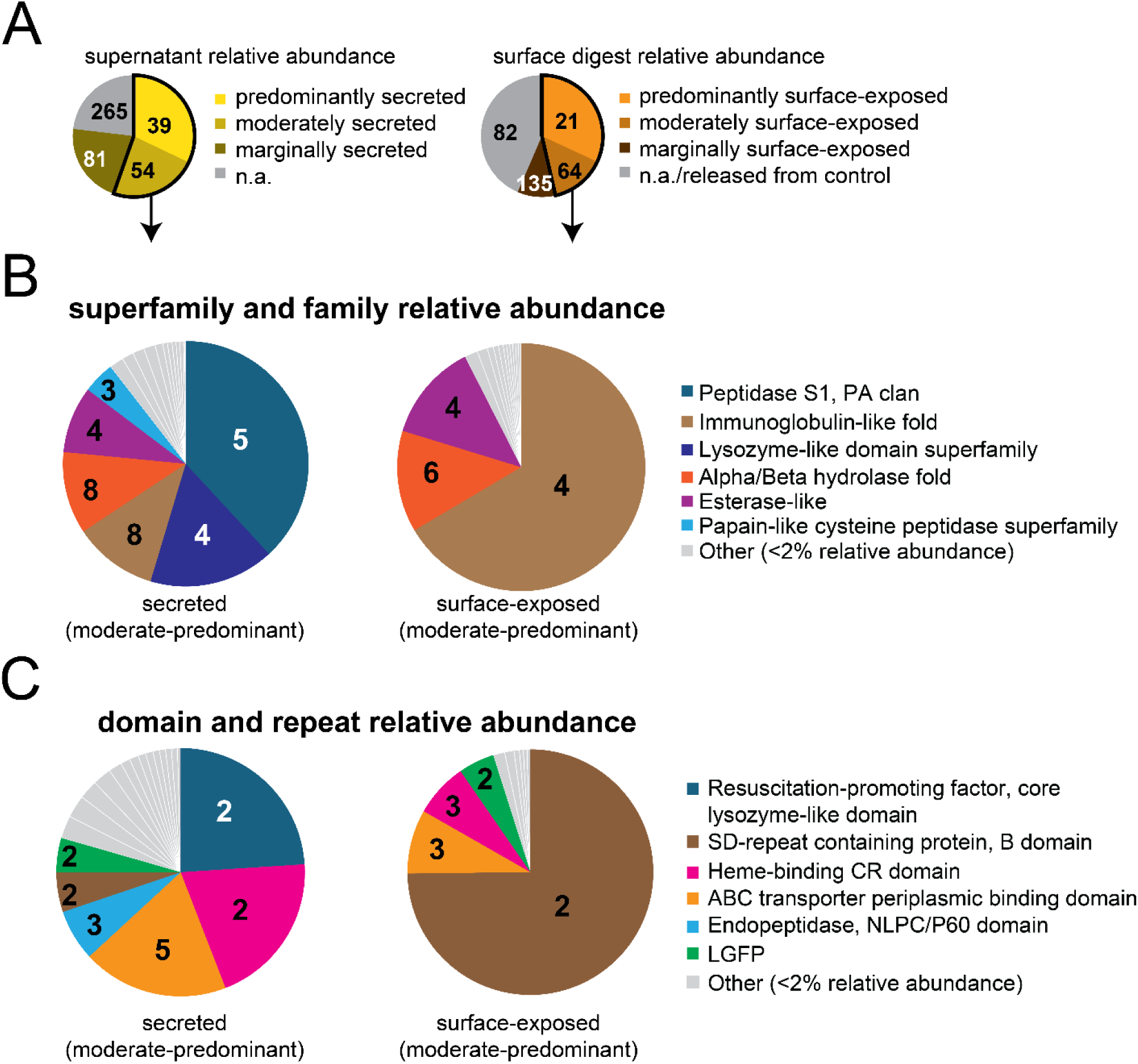
Highly abundant protein families and functional domains that are preferentially secreted (left) or surface-exposed (right) by *C. diphtheriae* under iron-limiting conditions. (A) relative abundances of all proteins quantified in the supernatant (left) or surface proteolysis (right) based on their preference for secretion or surface-exposure, respectively. Predominantly, moderately, and marginally secreted proteins are generally defined as having >90%, >50-90%, and >30-50% secretion (Fig. 2A), while those that are “not applicable (n.a.)” have ≤30% secretion or do not have a statistically significant higher abundance in the supernatant relative to the cytoplasmic fraction (p-value < 0.05 as determined by a t-test). Predominantly, moderately, and marginally surface-exposed proteins have >9.8%, >2.0-9.8% or >0.1-2.0% of their total whole-cell culture-normalized abundance liberated from the 15 minute protolysis, while those with ≤0.1% or those without a canonical secretion signal (SignalP6.0^49^) and released from the experimental control were classified as “not-applicable (n.a.)”. (B) Predominantly and moderately secreted (left) or surface-exposed (right) protein abundances summed according to shared superfamilies and families (as defined by InterproScan^63^, which are shared common evolutionarily structural folds, sequences of functions), or (C) functional domains and sequence repeats (as defined by InterproScan^63^). Numbers within the pie charts specify the number of proteins giving rise to that abundance. Slices in the pie charts in (B) and (C) were only included if they were found with at least two proteins and were non-redundant (did not consist of the same proteins). Grey slices outlined in white show the abundance distribution of other less abundant groupings.

### Exoproteome and surfaceome changes in response to iron-restriction and growth on human Hb

During an infection, *C. diphtheriae* utilizes surface-exposed and secreted virulence factors to overcome iron limitation. To evaluate the impact of iron levels and source on the exoproteome and surfaceome we conducted a differential expression analysis. *C. diphtheriae* was grown to log-phase in three conditions that varied in either the amount or source of iron that was present and include: (a) high-iron (condition 2a), (b) Fe-restricted + FeCl_3_ (condition 2b), and (c) Fe-restricted + Hb (conditions #2c) conditions. We initially examined the largest and most significant changes that occur in the exoproteome and surfaceome as a result of iron restriction (referred to as “large” changes; ≥10-fold and ≤0.01 p-value) (**Fig. 5**). Iron-restriction with Hb and FeCl_3_ relative to the high-iron condition showed large increases in secreted proteins (11 and 6, respectively) (**Fig. 5A, B**) and surface-exposed proteins (10 and 9, respectively) (**Fig. 5D, E**). The largest effects on both proteomes occurred when cells were cultured in iron-restricted media that contained Hb as the iron source as compared to cells grown in high-iron. Specifically, large increases were observed for secreted DT, and a combination of secreted and surface-exposed iron-siderophore/heme ABC transporter solute binding components (Irp6A, FrgD, HmuT), species-conserved heme-iron acquisition system components (HtaA, HtaB), strain-specific heme-binding proteins (ChtA, ChtB, ChtC), and a hemoglobin binding protein (HbpA) (**Fig. 5A, 5D**). Only two proteins’ abundance largely decreased when cells were cultured in low-iron media; the uncharacterized DUF5926 protein liberated from surface proteolysis (**Fig. 5E**), and ferritin (Ftn), a cytoplasmic iron-storage protein that is also found in the supernatant (**Fig. 5B**). No large changes were observed between iron-restricted conditions (**Fig. 5C, F**). A complete list summarizing the iron-dependence on the secretion and surface-exposure of these proteins is in **Table S4**.

**Figure 5:**
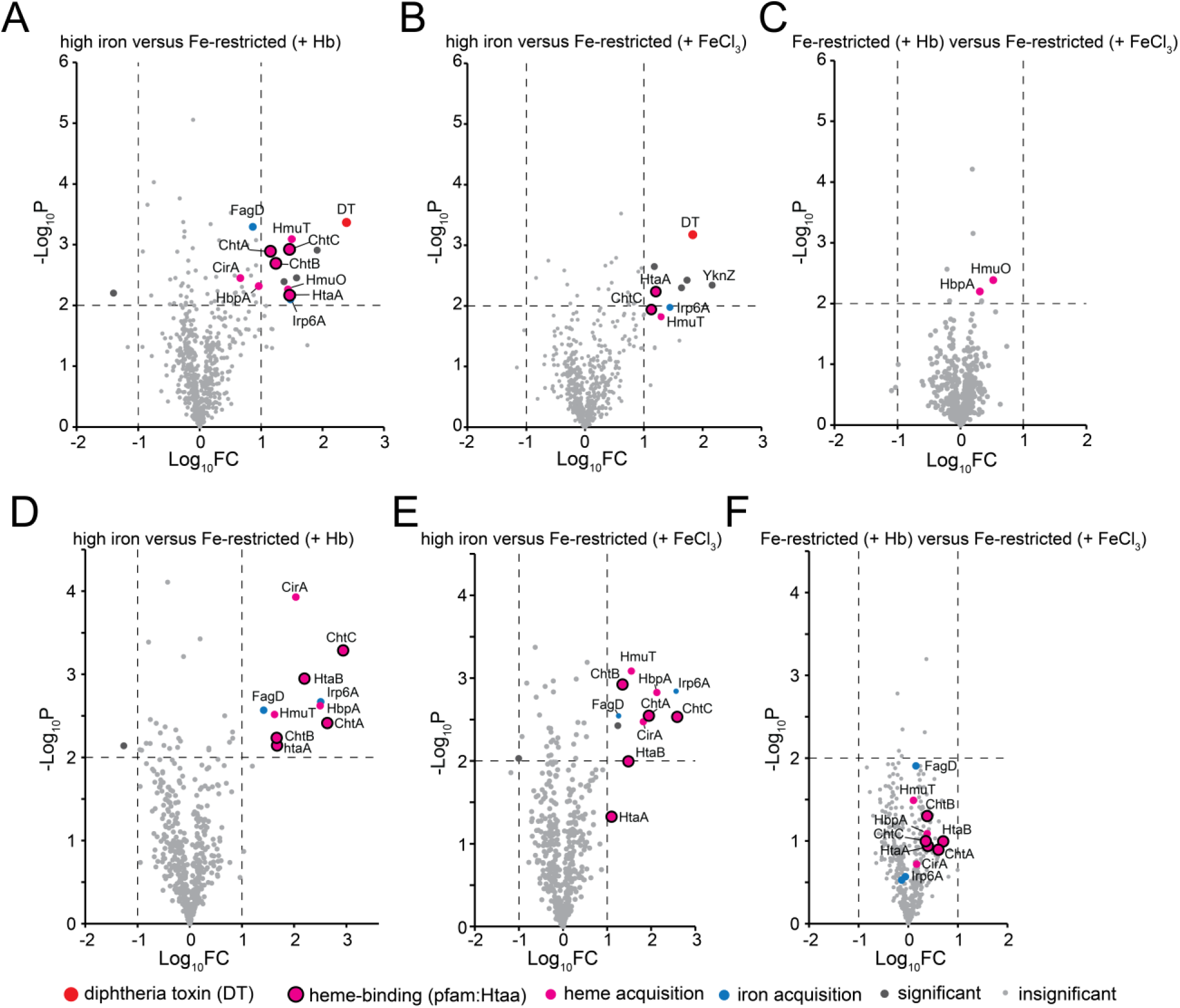
Highly significant and large alterations in the exoproteome and surfaceome caused by iron abundance and source. Volcano plots from protein MaxLFQ values^64^ were created using stringent significance cut-offs (p-value < 0.01) and fold-change requirements (>10-fold) to highlight only the most “drastic” changes between iron conditions for (A-C) supernatant fractions or (D-F) from surface-proteolyzed fractions. Volcano plots show differences between high-iron and (A, D) iron-restricted conditions (mPGT + 10µM Fe-chelator EDDHA) with 0.3 µM Hb supplementation on a heme-iron iron basis, or (B, E) iron-restricted conditions with 10 µM FeCl_3_ supplementation (equivalent concentration to Fe-chelator). (C, F) Differences between iron-restricted conditions with either Hb or FeCl_3_ supplementation. The diphtheria toxin (DT) is colored red, putative components of heme-uptake are pink, those containing a heme-binding CR domain (pfam:Htaa) are outlined in black, and putative iron and iron-siderophore acquisition components are colored blue if they have either a p-value of <0.01 or fold-change >10. All other proteins with drastic changes in secretion or surface-exposure are dark grey.

To see what other secreted and surface-exposed proteins shared iron-dependent expression profiles with the prominent virulence factors (DT and heme-acquisition system components), for each fraction we compared abundance differences across all three iron conditions (high-iron versus iron-restricted + Hb versus iron-restricted + FeCl_3_ conditions). An analysis of variance (ANOVA) was performed to find significant differences in protein abundance caused by at least one iron-condition, and all significantly altered proteins were clustered based on their expression profile. Major differences are summarized using a heatmap for the exoproteome (**Fig. 6A**) and surfaceome (**Fig. 6B**), and additional comparative data is presented in **Table S4**. This analysis revealed that when cells are deprived of iron, a similar set of proteins increase in abundance in the secreted and surface fractions when either FeCl_3_ or Hb is provided as the sole source of iron, indicated by red clusters in **Figure 6**. Additionally, our analysis reveals that DT is secreted more in both types of iron-restricted media, but the highest level of secretion occurs when Hb is provided as the sole iron source. Interestingly, hemoglobin-binding protein, HbpA, and corynemycolic acid transferase C (cMytC, also known as surface-layer protein A, SlpA), followed the same secretion profile (**Fig 6A**), and are among the most abundant and secreted proteins as determined in our low-iron preferential localization experiments (**Fig. 2**, **Table S2**). In the surface-digests, heme-binding HtaB, ChtA, and ChtB proteins were shown to follow the same trends (**Fig. 6B**). Fewer extracellular proteins had the opposite trend, where they were significantly higher under high-iron relative to either of the iron-restricted conditions (**Fig. 6**, **Table S4**). Notably, only one protein in the exoproteome (RipA) and 4 proteins from the surface digest (PiuB, DIP1481, DIP1632, DIP2036) were higher exclusively when Hb was the sole iron source (**Fig. 6**, **Table S4**).

**Figure 6:**
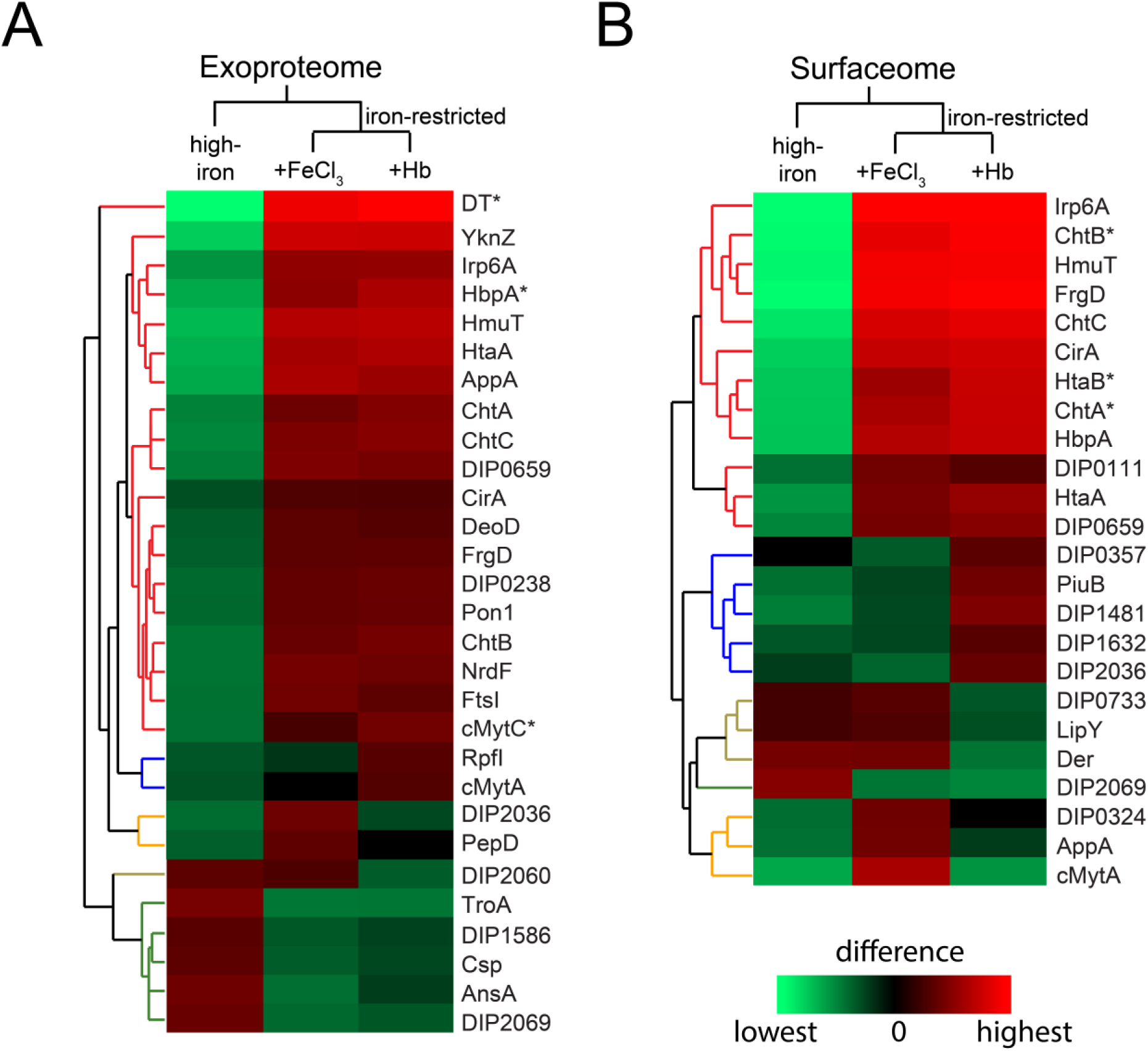
Significant iron-dependent alterations in the exoproteome and surfaceome. An ANOVA analysis was ran on the MaxLFQ values^64^ obtained from (A) supernatant and (B) surface-proteolyzed fractions to determine statistically significant (Benjamini-Hochberg fdr-adjusted p-value cut-off < 0.05; S0 of 0.0) changes caused by at least one iron condition. The heat map is colored based on significant differences determined from a Tukey’s post-hoc test following the ANOVA significance test, with red higher than green, and pure black indicating no significant difference. Proteins shown contain either a secretion signal or transmembrane helix as determined from SignalP 6.0^49^, CW-Pred^52^, and DeepTMHMM^51^. Clustering of proteins from each fraction were performed in Perseus.^65^ Clusters are shown in different colors on the dendrogram and represent groups of proteins with similar expression profiles. Clusters marked in red show the strongest increases under iron-restriction, and those significantly higher with Hb as the sole iron are marked by an asterisk. Blue clusters show proteins higher in Hb-fed iron-restricted cultures. For the exact significant relationships between each iron condition for the proteins shown here, refer to **Table S4**.

## DISCUSSION

*C. diphtheriae* virulence is strain-dependent,^66^ with close relatives of a *tox*-bearing *gravis* biotype responsible for the large-scale 1990s diphtheria pandemic encoding the greatest number of virulence factors.^67^ To gain insight into the bacterial machinery responsible for host colonization, we used cell fractionation and quantitative proteomics to define the exoproteome and surfaceome of *C. diphtheriae* strain 1737, which is widely used in published studies and a close relative of the pandemic strain.^4,58^ Initially, we evaluated this microbe’s proteome in low-iron conditions that are known to stimulate toxin production and other virulence factors (condition 1).^10,30,32,37,68^ We employed a label-free LC-MS/MS method to quantify each protein’s abundance in four distinct subcellular locations: secreted, cell envelope, surfaced-exposed, and cytoplasmic fractions. In this analysis, a protein’s abundance is normalized with respect to its total amount in the cell, thereby generating its “localization profile” that unambiguously reports on its preferred location (**Fig. 1**, **Table S1**). This approach is particularly useful for defining a protein’s level of secretion (**Fig. 2**) and surface-exposure (**Fig. 3**), and is to the best of our knowledge the first time it has been applied to a bacterium.^62^ Combined with a bioinformatics analysis the localization profiles identify functionally important and abundant secreted and surface-exposed proteins in *C. diphtheriae* (**Fig. 4**). Furthermore, we determined how the exoproteome and surfaceome changes in response to iron depletion (**Fig. 5**, **Fig. 6**), an established environmental signal that during infections triggers proteome changes that facilitate host colonization. Together, this analysis provides insight into putative virulence factors and their spatial localization that may be important for host-pathogen interactions during an infection (**Fig. 7**).

**Figure 7:**
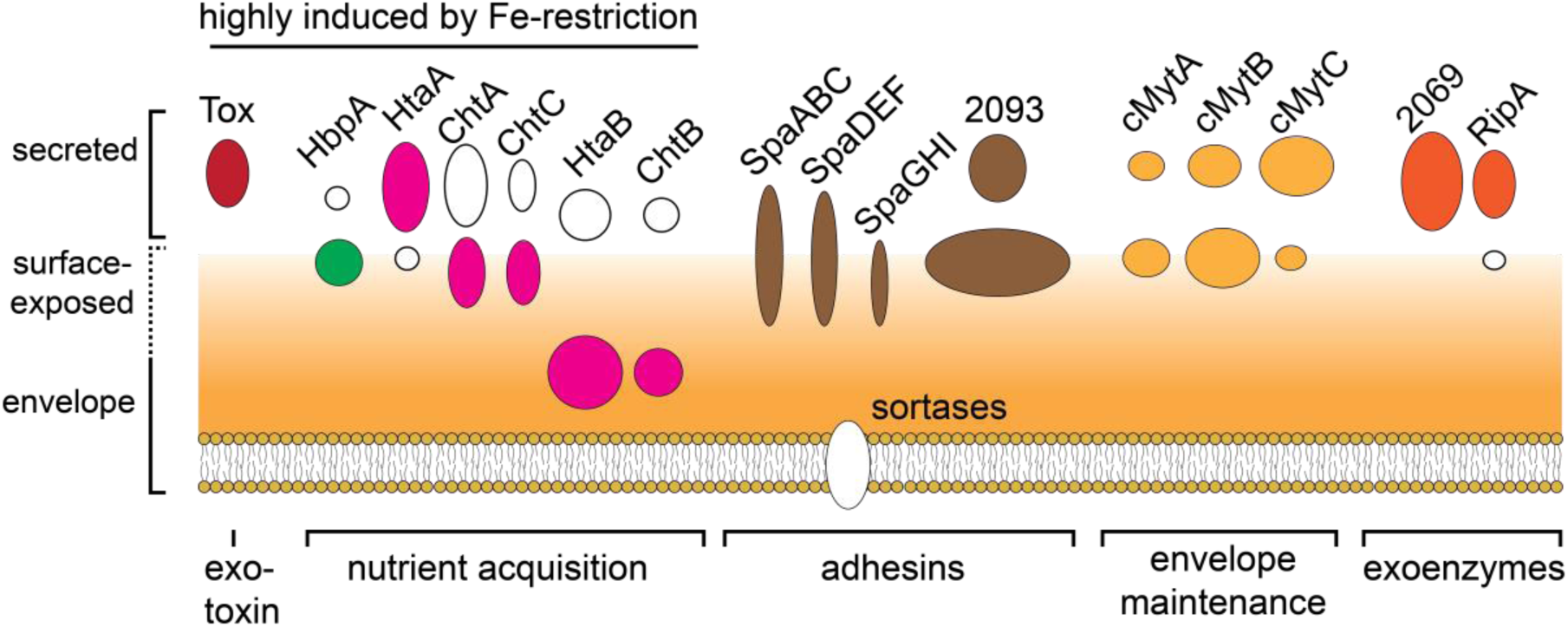
Predicted primary locations of select abundant proteins identified in our surface and secreted analysis. Proteins that are further from the surface have a higher preference for secretion, and proteins closer to the surface have a higher preference for surface-exposure. Proteins in the plasma membrane are shown for illustration purposes if identified in this study but were left uncolored as they were not evaluated for localization. Proteins that were drastically differentially secreted or surface-exposed in at least one iron-restricted condition (with Hb or FeCl_3_ supplementation) relative to the high-iron condition are shown on the left (Fig. 5). For proteins that are illustrated with two shapes, the larger shape indicates the primary localization (that is, the exoproteome or surfaceome depending on the higher localization preference where predominant > moderate > marginal); the smaller shape indicates the secondary location if it has moderate or higher secretion; if shapes are the same size, they are tied in location preference. The diphtheria toxin (DT) is shown in red. Heme-uptake proteins that encode two domains are shown as ovals (HtaA, ChtA, ChtC) and those with one domain as circles (HtaB, ChtB, HbpA); those encoding the canonical heme-binding CR domain are in pink while HbpA which encodes a domain involved in hemoglobin binding is shown in green. Putative adhesins are shown in brown (SpaABC, SpaDEF, SpaGHI, DIP2093), with a representative sortase responsible for cell wall anchoring shown in the membrane and left uncolored as plasma membrane proteins were not evaluated for localization. Corynemycolic acid transferases (cMytA, cMytB, cMytC) are shown in pale yellow. Select abundant exoenzymes from the trypsin-like protease (Clan PA) (DIP2069) and NlpC/P60 cysteine-endopeptidase (Clan CA) (RipA) indicated in orange, with a small dot indicating their identification on the surface albeit at small abundances. Proteins that were highly differentially expressed in one or both Fe-restricted conditions relative to the iron-replete condition (>10-fold increase, p-value < 0.01) are indicated.

### Exoenzymes are highly abundant and preferentially secreted in low-iron environments

Secreted proteins in pathogenic bacteria perform a variety of functions including: degrading host cell connective tissues and macromolecules for nutrients,^24,69^ post-translational proteolytic processing,^25,70^ quorum sensing,^71^ and nutrient acquisition^27^ among others. When *C. diphtheriae* is grown to mid log-phase in low-iron conditions (condition 1), our cellular fractionation experiments indicate that approximately one third of the total protein produced by the microbe is secreted (**Fig. 1B**). A total of 174 distinct proteins were reliably identified as being preferentially secreted (predominantly, moderately, and marginally secreted) based on their localization profiles (**Fig. 2A**) (**Table S1**). **Figure 2B** shows the most abundant proteins in the supernatant under low-iron conditions, of which 8 are predominantly secreted, 5 are moderately secreted, and 1 is partially secreted. The most abundant secreted protein is the serine protease, DIP2069 (**Fig. 7**), which constitutes almost 10% of the exoproteome and has a strong preference for secretion, with more than 99% of its total abundance found in the supernatant. Homologues of DIP2069 are highly abundant in the secreted proteomes of other *Corynebacterium* species.^72^ While its function is not known, DIP2069 belongs to the peptidase S1 (clan PA) superfamily; three other predominantly secreted proteases are also members of this family (DIP0743, DIP0964, DIP0350). Clan PA contains both serine and cysteine proteases that in other microbes participate in a range of processes, including nutrient acquisition, immune response suppression, tissue penetration, and posttranslational proteolytic processing of secreted virulence factors^21,70^. All these putative proteases consist of an N-terminal foldase domain that is essential for proper folding and inhibition of its catalytic C-terminal protease domain. The exoproteome also contains abundant proteases from clan CA, also known as NlpC/P60 cysteine endopeptidases (**Fig**. **4B**, **C**). Members of this clan possess a papain-like protein fold and have gained attention as critical virulence factors in a a variety of Actinobacterial species, including *M. tuberculosis*.^73^ Secreted clan CA enzymes include RipA, RipC, and RipC’ (**Fig. 7**). These proteins have homologues in other bacterial species such as *C. glutamicum* (Cg1735, Cg2401, Cg2401 respectively) and *M. tuberculosis* (Rv1477, Rv2190c, there is no homologue for RipC’).^74^ RipA may be involved in establishing the surface proteome for infection as genetic knockouts in *C. diphtheriae* ISS3319 resulted in attenuated adhesion to epithelial cells, reduced invasion in a *C. elegans* model, and a rearranged cell surface.^19^ RipA appears to be preferentially upregulated in iron-limited conditions that contain Hb (**Fig. 6**, **Table S4**), raising the possibility that it may also be important during an infection.

### Adhesins, membrane maintenance and iron acquisition proteins are the most abundant on the bacterial surface under low-iron conditions

The results of protease shaving experiments on iron-restricted cells reveal that some of the most abundant proteins on the cell surface have functions in microbial adhesion, outer mycolic acid membrane maintenance, and iron acquisition (**Fig. 3C**, **Fig. 4**). DIP2093 is the most abundant protein on the cell surface in these conditions and has been shown to bind collagen, and to promote bacterial adherence and invasion of host epithelial cells (**Fig. 3C, Fig. 7**).^75^ Another highly abundant and preferentially located surface-exposed adhesin is DIP2062 (**Fig. 3C**), which contains adhesive SdrB repeat domains. An analysis of the primary sequences of surface displayed proteins using the program CW-pred^52^ suggests that many of the surface adhesins are covalently attached to the underlying cell wall by sortases, which recognize a C-terminal cell wall sorting signal motif.^76^ The genome of *C. diphtheriae* encodes 16 proteins that are covalently assembled pili or attached to the cell wall by six distinct sortase cysteine transpeptidase enzymes (SrtA to SrtF).^76^ A total of 14 sortase substrates were identified by MS, 10 of which were found on the cell surface through our surface shaving experiment. These proteins include the aforementioned highly abundant DIP2062 and DIP2093 proteins, and 3 types of pili: SpaA-pili (formed from SpaA, SpaB, SpaC pilins), SpaD-pili (formed from SpaD, SpaE, SpaF pilins), and SpaH-pili (formed from SpaG, SpaH, and SpaI pilins) (**Fig. 7**). All the pilin components within these pili are sortase substrates and were identified by MS, with the exception of SpaI. Interestingly, the localization profiles of the pilins are unique as they exhibit moderate to predominant surface exposure, while they are also abundant in the supernatant. Their presence in the secreted fraction could be explained by their shearing from cells during agitated growth. Based on the sum of their abundances in the MS data, the SpaA- and SpaD-pili are ∼8-fold more abundant than SpaH-pili when cells are grown in iron-limiting conditions (**Table S1**). Interestingly, the compositions of the SpaA- and SpaD-pili appears to vary, as the tip pilin in the SpaA-pilus (SpaC) is ∼6 fold more abundant than its shaft pilin (SpaA), whereas the analogous components in the SpaD-pilus (SpaD (shaft) and SpaF (tip) pilins) are equally abundant (**Table S1**). Collectively, our results indicate that during log-phase growth in low-iron conditions, *C. diphtheriae* displays covalently attached surface adhesins such as DIP2093, DIP2062, and SpaA/SpaH/SpaF pili that could function to promote microbial adhesion to host tissues.

Many proteins exhibit dual localization profiles as they are both secreted and located on the cell surface in iron limiting conditions. Three corynemycolic acid transferase proteins exhibit this phenotype are very abundant: cMytA, cMytB, and cMytC (**Fig. 7**). Based on sequence homology they are likely part of the mycobacterial Ag85 complex whose components esterify arabinogalactan and trehalose to mycolic acids to form the inner and outer leaflet of the mycolic acid layer, respectively.^77^ ^78^ Interestingly, cMytC was slightly more abundant in the secretome than cMytA and cMytB, despite its presumed role in modifying the inner leaflet. We speculate that its presence in the secreted fraction may result from the production of outer myco-membrane vesicles (MMVs), which have previously been observed in the closely-related bacterium *C. glutamicum* when it is cultured in low-iron media.^79,80^ Interestingly, cMytA is also significantly higher in the supernatants of iron-restricted cultures grown in the presence of Hb, which could be explained through its presence in mycomembrane vesicles, although further experimental confirmation would be required to substantiate this. How these corynemycolic acid transferases are localized to the surface remains unclear, but it could be mediated by posttranslational modification of cMytB and cMytC with mycolic acid as they contain the appropriate “LSS” sequence motif for this alteration,^81^ while in cMytA it could result from non-covalent interactions with the envelope that occur via its LGFP domain.^82^ These proteins are also prevalent in the supernatant, suggesting that these interactions with the mycolic acid layer are weak. Several proteins involved in heme-uptake are also secreted and present on the cell surface when iron is limiting and are discussed below.

### Highly abundant surface-exposed and secreted proteins with unknown functions

Actinobacteria are unique, as they possess an outer mycolic acid layer whose width and lipid composition varies between *Corynebacterium*, *Mycobacterium*, and *Norcardia* species. The protein machinery embedded in this unique structure is generally not well characterized, and primarily only low-molecular weight pore-forming proteins have been identified (typically ∼10-15 kDa). Interestingly, our results indicate that two non-polar proteins are likely located in the mycolic acid and upregulated when iron is restricted, DIP0659 and CirA. DIP0659 has not been characterized and is the second most abundant protein in the surfaceome. It is relatively large (41 kDa) and especially upregulated under iron-restriction (**Fig. 6**, **Table S4**). The function of CirA is also not known, but it is expressed from the same operon that encodes the ChtC hemoprotein raising the intriguing possibility that it may facilitate heme passage across the mycolic acid layer.^10,30^ There are also several abundant and predominantly secreted proteins whose functions are not well understood. This includes DIP0013, which is a small 16.6 kDa protein that contains a Rib-long domain implicated in protective immunity.^83^ Another highly abundant secreted protein, DIP2014, shows dual localization between the exoproteome and cell envelope or surface. It is a BigA-like protein which has a sequence homolog in pathogenic *Brucella abortus* that elicits cytoskeleton rearrangements in HeLa cells that may be critical for host colonization.^84^

### Insight into how components in the heme-uptake system are positioned

*C. diphtheriae*’s heme acquisition system contains at least five hemoproteins (HtaA, HtaB, ChtA, ChtB, ChtC) that bind heme via conserved CR domains,^10,29,30^ an ABC transporter (HmuTUV) that imports heme across the membrane,^85–87^ a heme oxygenase (HmuO),^88^ and HbpA which binds to Hb and the Hb-haptoglobin (Hb-Hp) complex.^31^ **Figure S6** shows the localization profiles and relative abundance of these components. While all CR-containing proteins and HbpA contain a C-terminal transmembrane helix suggesting they may be embedded in the membrane, they nevertheless exhibit very distinct localization profiles (**Fig. 7**).^70^ In particular, HtaA is primarily secreted suggesting that it functions as hemophore that scavenges heme from the cell’s surroundings (**Fig. 7**). In contrast, the paralogous ChtA and ChtC hemoproteins are located on the microbial surface based on our proteolytic shaving experiments, while fractionation experiments indicate that a significant amount of ChtA is also secreted (**Fig. 7**). The Hb binding HbpA receptor also primarily resides on the surface suggesting that it may work in concert with the ChtA/ChtC paralogues to remove heme from Hb at this site (**Fig. 7**).^27^ The remaining CR containing hemoproteins in the system, HtaB and ChtB, are presumably located near the extracellular cytoplasmic membrane because they primarily localize to the envelope fraction and they are only minimally susceptibility to proteolysis in the shaving experiments (**Fig. 7**). As expected, the HmuO oxygenase resides within the cytoplasm, consistent with it degrading heme after it is imported into the cell. Interestingly, the HmuT heme binding protein is the most abundant heme-uptake component, and a majority of its abundance comes from the supernatant. This location is surprising, as HmuT is predicted to be lipid-modified and generally thought to be positioned on the extracellular cytoplasmic membrane where it functions as the solute binding component of the ABC HmuTUV transporter that pumps heme into the cytoplasm. Notably, in addition to HmuT several other solute binding components of ABC transporters are also present in the supernatant at high relative abundance even though they are presumably lipidated (FrgD, DIP1570, CtaC). As both HmuT and HtaA are prevalent in the secreted fraction even though their primary sequences contain membrane anchoring elements (a lipid modification site and transmembrane helix, respectively), our results suggest that they may only be weakly bound to the plasma membrane enabling their facile detachment. It is also possible that these proteins are released from the cell as a result of proteolytic processing, as observed for several hemophore-like proteins in the oral pathogen *P. gingivalis*.^70^ While our analysis clearly reveals the most abundant location of each component in the uptake system (“primary location”, **Fig. S6**), many of these proteins appear to be present in multiple locations (e.g. embedded in the cell envelope, located on the surface and secreted). This suggests that depending upon its location, each protein might play multiple roles in heme-uptake, from scavenging heme in the environment that surrounds the microbe to facilitating its transfer across the expanse of the cell wall. However, it cannot be ruled out that this distribution is caused by fractional bleeding during the fractionation process.

### Hb in the growth media accentuates the effects of iron-restriction

To gain insight into how *C. diphtheriae* combats the human nutritional immunity response,^32,89^ we determined how its exoproteome and surfaceome changed when it is grown in iron-restricted media that contained either Hb or FeCl_3_ as a sole source of iron. Based on previously reported transcriptomic studies several proteins should exhibit iron-dependent abundance changes because the expression of their genes is controlled by the iron-responsive DtxR repressor and/or two heme/hemoglobin-dependent two-component response systems – HrrSA and ChrSA. The DtxR regulon is well described for *C. diphtheriae*^32,37,38,90^ and controls the up- and down-regulation of a variety of genes encoding proteins involved iron metabolism, acquisition, and storage, as well as non-iron related virulence factors (e.g. DT).^37^ Proteins encoded by genes within the ChrSA and HrrSA regulons maintain heme homeostasis by mediating its synthesis, acquisition and export.^40^ For example, in response to elevated levels of heme the ChrSA/HrrSA systems increase the expression of genes encoding the cytoplasmic heme-degrading protein HmuO, the heme-exporting HrtAB transporter,^91^ and they repress the expression of genes that encode enzymes involved in heme biosynthesis.^92^ Since a main source of iron in the human body is heme, it is not surprising that many genes are co-regulated by DtxR and the HrrSA/ChrSA systems. For instance, the *htaAB* and *chtAB* operons involved in heme-iron acquisition are regulated by DtxR and the HrrSA/ChrSA systems.

We hypothesized that when cells are grown in iron-restricted growth media, the presence of heme/hemoglobin may act to boost the amount of select iron-dependent genes whose protein products promote heme acquisition and metabolism, and possibly other virulence factors that optimize these processes. To investigate this issue, we compared the proteomes of cells cultured in three distinct types of growth media: a high-iron media (condition 2a) versus iron-restricted media in which either Hb (condition 2b) or free iron was provided as the sole source of iron (condition 2c). Interestingly, a rigorous 3-way ANOVA analysis of this data reveals statistically significant iron source-dependent effects on protein abundance (**Fig. 6**). In particular, many heme-binding proteins whose genes are regulated by both HrtSA/ChrSA and DtxR are upregulated in iron-restricted conditions when either nutrient source is present, but their abundance is further increased when cells are grown in Hb. Increased expression when Hb is the iron source is observed for HbpA (the Hb receptor) and several heme-binding CR domain-containing proteins (HtaB, ChtA, and ChtB). Moreover, several proteins seemingly unrelated to heme-iron acquisition exhibit similar Hb-enhanced expression profiles. In the exoproteome, the most pronounced effect is observed by the diphtheria exotoxin (DT), which is cytotoxic and causes the high mortality associated with diphtheria.^1,66^ Additionally, levels of corynemycolic acid transferase C (cMytC), also known as S-layer protein (SlpA) are enhanced when Hb is present, suggesting that broad changes in the microbial surface occur that could aid in adhesion, immune system evasion, and possibly nutrient acquisition^93,94^. Increased secretion of RipA (DIP1821, also known as RpfI) in Hb-fed cultures also suggests it is implicated in virulence, as RipA has been shown to be critical for adhesion and internalization into epithelial cells (**Fig. 6A**, **Table S4**).^19^ On the cell surface, Hb-enhanced surface-exposure is observed for PiuB; this protein encodes a PepSY domain that has been suggested to regulate protease activity in the local environment.^95^ Thus, PiuB could be responsible for a broad range of effects caused by proteolytic processing of surface proteins and cell wall hydrolases, which are often regulated through cleavage events.^73^ Interestingly, three relatively uncharacterized proteins are also increased at the cell surface exclusively Hb-enhanced. Of these, DIP1481 contains a domain of unknown function (DUF), DUF4439, and belongs to the Ferritin-like superfamily, members of which not only bind iron, but also other free metal and metal-ligated species (e.g. heme, covalbumin).^96^ Thus, DIP1481 may be involved in heme-iron storage. Less can be postulated about the roles of a DUF3043-containing DIP1632, and DIP2036 which has no close sequence homologues by BLASTP search (DIP2036). However, based on their surface-exposure profiles, they serve as good candidates for future microbial and biochemical studies for their impact on pathogenicity.

In conclusion, we applied a label-free quantitative proteomics workflow to identify with high confidence secreted and surface-exposed proteins in *C. diphtheriae* when it is grown in low-iron conditions that contain either Hb or FeCl_3_. By assessing differential secretion and surface-exposure in multiple conditions we determined the effects of iron abundance and iron-source on the proteome. The majority of the exoproteome contains exoproteases and hydrolases, while the surfaceome is primarily comprised of adhesins, with DIP2093 being 4 times more abundant than any other protein on the cell surface. Heme-binding proteins are also highly abundant in both locations, with HtaA primarily located in the secreted fraction where it may function as hemophore to scavenge heme from the microbe’s surrounding, whereas the heme-binding ChtA and ChtC proteins are positioned on the cell surface in virulent bacterial strains where they may function to receive heme from HtaA. In addition, the presence of Hb in iron-restricted media selectively enhances the abundance of several proteins that may facilitate host colonization, including the surface displayed DIP0659. Collectively, these findings provide new insight into the composition of the exoproteome and surfaceome in *C. diphtheriae* and related actinobacteria that cause disease, and could thus could facilitate the discovery of new drugs to treat antibiotic-resistant bacterial infections.

## ACKNOWLEDGMENTS

We thank members of the Clubb and Loo laboratories for useful discussions. This work was funded by grants from the National Institutes of Health R01AI161828 (to RTC and JAL) and R35GM145286 (to JAL). A.K.G. acknowledges support from the NIH National Institute of Dental & Craniofacial Research (T90DE030860). We thank Dr. Michael Schmitt for providing *C. diphtheriae* strain 1737.

## REFERENCES

(1) Corynebacterium Diphtheriae and Related Toxigenic Species; Burkovski, A., Ed.; Springer Netherlands: Dordrecht, 2014. 10.1007/978-94-007-7624-1.

(2) Sharma, N. C.; Efstratiou, A.; Mokrousov, I.; Mutreja, A.; Das, B.; Ramamurthy, T. Diphtheria. Nat. Rev. Dis. Primer 2019, 5 (1), 81. 10.1038/s41572-019-0131-y.

(3) Will, R. C.; Ramamurthy, T.; Sharma, N. C.; Veeraraghavan, B.; Sangal, L.; Haldar, P.; Pragasam, A. K.; Vasudevan, K.; Kumar, D.; Das, B.; Heinz, E.; Melnikov, V.; Baker, S.; Sangal, V.; Dougan, G.; Mutreja, A. Spatiotemporal Persistence of Multiple, Diverse Clades and Toxins of Corynebacterium Diphtheriae. Nat. Commun. 2021, 12 (1), 1500. 10.1038/s41467-021-21870-5.

(4) Popovic, T.; Kombarova, S. Y.; Reeves, M. W.; Nakao, H.; Mazurova, I. K.; Wharton, M.; Wachsmuth, I. K.; Wenger, J. D. Molecular Epidemiology of Diphtheria in Russia, 1985--1994. J. Infect. Dis. 1996, 174 (5), 1064–1072. 10.1093/infdis/174.5.1064.

(5) Goodall, E. C. A.; Azevedo Antunes, C.; Möller, J.; Sangal, V.; Torres, V. V. L.; Gray, J.; Cunningham, A. F.; Hoskisson, P. A.; Burkovski, A.; Henderson, I. R. A Multiomic Approach to Defining the Essential Genome of the Globally Important Pathogen Corynebacterium Diphtheriae. PLOS Genet. 2023, 19 (4), e1010737. 10.1371/journal.pgen.1010737.

(6) Hennart, M.; Panunzi, L. G.; Rodrigues, C.; Gaday, Q.; Baines, S. L.; Barros-Pinkelnig, M.; Carmi-Leroy, A.; Dazas, M.; Wehenkel, A. M.; Didelot, X.; Toubiana, J.; Badell, E.; Brisse, S. Population Genomics and Antimicrobial Resistance in Corynebacterium Diphtheriae. Genome Med. 2020, 12 (1), 107. 10.1186/s13073-020-00805-7.

(7) Trost, E.; Blom, J.; de Castro Soares, S.; Huang, I.-H.; Al-Dilaimi, A.; Schroder, J.; Jaenicke, S.; Dorella, F. A.; Rocha, F. S.; Miyoshi, A.; Azevedo, V.; Schneider, M. P.; Silva, A.; Camello, T. C.; Sabbadini, P. S.; Santos, C. S.; Santos, L. S.; Hirata, R.; Mattos-Guaraldi, A. L.; Efstratiou, A.; Schmitt, M. P.; Ton-That, H.; Tauch, A. Pangenomic Study of Corynebacterium Diphtheriae That Provides Insights into the Genomic Diversity of Pathogenic Isolates from Cases of Classical Diphtheria, Endocarditis, and Pneumonia. J. Bacteriol. 2012, 194 (12), 3199–3215. 10.1128/JB.00183-12.

(8) Broadway, M. M.; Rogers, E. A.; Chang, C.; Huang, I.-H.; Dwivedi, P.; Yildirim, S.; Schmitt, M. P.; Das, A.; Ton-That, H. Pilus Gene Pool Variation and the Virulence of Corynebacterium Diphtheriae Clinical Isolates during Infection of a Nematode. J. Bacteriol. 2013, 195 (16), 3774–3783. 10.1128/JB.00500-13.

(9) Schmitt, M. P. Corynebacterium Diphtheriae. In Iron Transport in Bacteria; Crosa, J. H., Mey, A. R., Payne, S. M., Eds.; ASM Press: Washington, DC, USA, 2014; pp 344–359. 10.1128/9781555816544.ch22.

(10) Allen, C. E.; Schmitt, M. P. Utilization of Host Iron Sources by Corynebacterium Diphtheriae: Multiple Hemoglobin-Binding Proteins Are Essential for the Use of Iron from the Hemoglobin-Haptoglobin Complex. J. Bacteriol. 2015, 197 (3), 553–562. 10.1128/JB.02413-14.

(11) Biology and Biotechnology of Actinobacteria; Wink, J., Mohammadipanah, F., Hamedi, J., Eds.; Springer International Publishing: Cham, 2017. 10.1007/978-3-319-60339-1.

(12) Uhde, K. B.; Pathak, S.; McCullum Jr, I.; Jannat-Khah, D. P.; Shadomy, S. V.; Dykewicz, C. A.; Clark, T. A.; Smith, T. L.; Brown, J. M. Antimicrobial-Resistant *Nocardia* Isolates, United States, 1995–2004. Clin. Infect. Dis. 2010, 51 (12), 1445–1448. 10.1086/657399.

(13) Bacterial Cell Walls and Membranes; Kuhn, A., Ed.; Subcellular Biochemistry; Springer International Publishing: Cham, 2019; Vol. 92. 10.1007/978-3-030-18768-2.

(14) Nguyen, L. Antibiotic Resistance Mechanisms in M. Tuberculosis: An Update. Arch. Toxicol. 2016, 90 (7), 1585–1604. 10.1007/s00204-016-1727-6.

(15) Kasik, J. E.; Peacham, L. Properties of β-Lactamases Produced by Three Species of Mycobacteria. Biochem. J. 1968, 107 (5), 675–682. 10.1042/bj1070675.

(16) Chambers, H. F.; Moreau, D.; Yajko, D.; Miick, C.; Wagner, C.; Hackbarth, C.; Kocagöz, S.; Rosenberg, E.; Hadley, W. K.; Nikaido, H. Can Penicillins and Other Beta-Lactam Antibiotics Be Used to Treat Tuberculosis? Antimicrob. Agents Chemother. 1995, 39 (12), 2620–2624. 10.1128/AAC.39.12.2620.

(17) Casadevall, A.; Pirofski, L. Virulence Factors and Their Mechanisms of Action: The View from a Damage–Response Framework. J. Water Health 2009, 7 (S1), S2–S18. 10.2166/wh.2009.036.

(18) Hansmeier, N.; Chao, T.-C.; Kalinowski, J.; Pühler, A.; Tauch, A. Mapping and Comprehensive Analysis of the Extracellular and Cell Surface Proteome of the Human Pathogen *Corynebacterium Diphtheriae*. PROTEOMICS 2006, 6 (8), 2465–2476. 10.1002/pmic.200500360.

(19) Ott, L.; Höller, M.; Gerlach, R. G.; Hensel, M.; Rheinlaender, J.; Schäffer, T. E.; Burkovski, A. Corynebacterium Diphtheriae Invasion-Associated Protein (DIP1281) Is Involved in Cell Surface Organization, Adhesion and Internalization in Epithelial Cells. BMC Microbiol. 2010, 10 (1), 2. 10.1186/1471-2180-10-2.

(20) Möller, J.; Nosratabadi, F.; Musella, L.; Hofmann, J.; Burkovski, A. Corynebacterium Diphtheriae Proteome Adaptation to Cell Culture Medium and Serum. Proteomes 2021, 9 (1), 14. 10.3390/proteomes9010014.

(21) Bittel, M.; Gastiger, S.; Amin, B.; Hofmann, J.; Burkovski, A. Surface and Extracellular Proteome of the Emerging Pathogen Corynebacterium Ulcerans. Proteomes 2018, 6 (2), 18. 10.3390/proteomes6020018.

(22) Möller, J.; Schorlemmer, S.; Hofmann, J.; Burkovski, A. Cellular and Extracellular Proteome of the Animal Pathogen Corynebacterium Silvaticum, a Close Relative of Zoonotic Corynebacterium Ulcerans and Corynebacterium Pseudotuberculosis. Proteomes 2020, 8 (3), 19. 10.3390/proteomes8030019.

(23) Silva, W. M.; Carvalho, R. D. D. O.; Dorella, F. A.; Folador, E. L.; Souza, G. H. M. F.; Pimenta, A. M. C.; Figueiredo, H. C. P.; Le Loir, Y.; Silva, A.; Azevedo, V. Quantitative Proteomic Analysis Reveals Changes in the Benchmark *Corynebacterium Pseudotuberculosis* Biovar Equi Exoproteome after Passage in a Murine Host. Front. Cell. Infect. Microbiol. 2017, 7, 325. 10.3389/fcimb.2017.00325.

(24) Malloy, J. L.; Veldhuizen, R. A. W.; Thibodeaux, B. A.; O’Callaghan, R. J.; Wright, J. R. *Pseudomonas Aeruginosa* Protease IV Degrades Surfactant Proteins and Inhibits Surfactant Host Defense and Biophysical Functions. Am. J. Physiol.-Lung Cell. Mol. Physiol. 2005, 288 (2), L409–L418. 10.1152/ajplung.00322.2004.

(25) Rigoulay, C.; Entenza, J. M.; Halpern, D.; Widmer, E.; Moreillon, P.; Poquet, I.; Gruss, A. Comparative Analysis of the Roles of HtrA-Like Surface Proteases in Two Virulent *Staphylococcus Aureus* Strains. Infect. Immun. 2005, 73 (1), 563–572. 10.1128/IAI.73.1.563-572.2005.

(26) Barocchi, M. A.; Ries, J.; Zogaj, X.; Hemsley, C.; Albiger, B.; Kanth, A.; Dahlberg, S.; Fernebro, J.; Moschioni, M.; Masignani, V.; Hultenby, K.; Taddei, A. R.; Beiter, K.; Wartha, F.; Von Euler, A.; Covacci, A.; Holden, D. W.; Normark, S.; Rappuoli, R.; Henriques-Normark, B. A Pneumococcal Pilus Influences Virulence and Host Inflammatory Responses. Proc. Natl. Acad. Sci. 2006, 103 (8), 2857–2862. 10.1073/pnas.0511017103.

(27) Choby, J. E.; Skaar, E. P. Heme Synthesis and Acquisition in Bacterial Pathogens. J. Mol. Biol. 2016, 428 (17), 3408–3428. 10.1016/j.jmb.2016.03.018.

(28) Antelo, G. T.; Vila, A. J.; Giedroc, D. P.; Capdevila, D. A. Molecular Evolution of Transition Metal Bioavailability at the Host–Pathogen Interface. Trends Microbiol. 2021, 29 (5), 441–457. 10.1016/j.tim.2020.08.001.

(29) Allen, C. E.; Schmitt, M. P. HtaA Is an Iron-Regulated Hemin Binding Protein Involved in the Utilization of Heme Iron in Corynebacterium Diphtheriae. J. Bacteriol. 2009, 191 (8), 2638–2648. 10.1128/JB.01784-08.

(30) Allen, C. E.; Burgos, J. M.; Schmitt, M. P. Analysis of Novel Iron-Regulated, Surface-Anchored Hemin-Binding Proteins in Corynebacterium Diphtheriae. J. Bacteriol. 2013, 195 (12), 2852–2863. 10.1128/JB.00244-13.

(31) Lyman, L. R.; Peng, E. D.; Schmitt, M. P. Corynebacterium Diphtheriae Iron-Regulated Surface Protein HbpA Is Involved in the Utilization of the Hemoglobin-Haptoglobin Complex as an Iron Source. J. Bacteriol. 2018, 200 (7), e00676–17, /jb/200/7/e00676-17.atom. 10.1128/JB.00676-17.

(32) Merchant, A. T.; Spatafora, G. A. A Role for the DtxR Family of Metalloregulators in Gram-Positive Pathogenesis. Mol. Oral Microbiol. 2014, 29 (1), 1–10. 10.1111/omi.12039.

(33) Bilitewski, U.; Blodgett, J. A. V.; Duhme-Klair, A.-K.; Dallavalle, S.; Laschat, S.; Routledge, A.; Schobert, R. Chemical and Biological Aspects of Nutritional Immunity-Perspectives for New Anti-Infectives That Target Iron Uptake Systems. Angew. Chem. Int. Ed. 2017, 56 (46), 14360–14382. 10.1002/anie.201701586.

(34) Pappenheimer, A. M.; Johnson, S. J. Studies in Diphtheria Toxin Production. I: The Effect of Iron and Copper. 1936.

(35) Cescau, S.; Cwerman, H.; Létoffé, S.; Delepelaire, P.; Wandersman, C.; Biville, F. Heme Acquisition by Hemophores. BioMetals 2007, 20 (3–4), 603–613. 10.1007/s10534-006-9050-y.

(36) Boyd, J.; Oza, M. N.; Murphy, J. R. Molecular Cloning and DNA Sequence Analysis of a Diphtheria Tox Iron-Dependent Regulatory Element (dtxR) from Corynebacterium Diphtheriae. Proc. Natl. Acad. Sci. 1990, 87 (15), 5968–5972. 10.1073/pnas.87.15.5968.

(37) Yellaboina, S.; Ranjan, S.; Chakhaiyar, P.; Hasnain, S.; Ranjan, A. Prediction of DtxR Regulon: Identification of Binding Sites and Operons Controlled by Diphtheria Toxin Repressor in Corynebacterium Diphtheriae. BMC Microbiol. 2004, 4 (1), 38. 10.1186/1471-2180-4-38.

(38) Wittchen, M.; Busche, T.; Gaspar, A. H.; Lee, J. H.; Ton-That, H.; Kalinowski, J.; Tauch, A. Transcriptome Sequencing of the Human Pathogen Corynebacterium Diphtheriae NCTC 13129 Provides Detailed Insights into Its Transcriptional Landscape and into DtxR-Mediated Transcriptional Regulation. BMC Genomics 2018, 19 (1), 82. 10.1186/s12864-018-4481-8.

(39) Möller, J.; Kraner, M.; Sonnewald, U.; Sangal, V.; Tittlbach, H.; Winkler, J.; Winkler, T. H.; Melnikov, V.; Lang, R.; Sing, A.; Mattos-Guaraldi, A. L.; Burkovski, A. Proteomics of Diphtheria Toxoid Vaccines Reveals Multiple Proteins That Are Immunogenic and May Contribute to Protection of Humans against Corynebacterium Diphtheriae. Vaccine 2019, 37 (23), 3061–3070. 10.1016/j.vaccine.2019.04.059.

(40) Burgos, J. M.; Schmitt, M. P. The ChrSA and HrrSA Two-Component Systems Are Required for Transcriptional Regulation of the *hemA* Promoter in Corynebacterium Diphtheriae. J. Bacteriol. 2016, 198 (18), 2419–2430. 10.1128/JB.00339-16.

(41) Keppel, M.; Piepenbreier, H.; Gätgens, C.; Fritz, G.; Frunzke, J. Toxic but Tasty – Temporal Dynamics and Network Architecture of Heme-responsive Two-component Signaling in *Corynebacterium Glutamicum*. Mol. Microbiol. 2019, 111 (5), 1367–1381. 10.1111/mmi.14226.

(42) Tai, S.-P. S.; Krafft, A. E.; Nootheti, P.; Holmes, R. K. Coordinate Regulation of Siderophore and Diphtheria Toxin Production by Iron in Corynebacterium Diphtheriae. Microb. Pathog. 1990, 9 (4), 267–273. 10.1016/0882-4010(90)90015-I.

(43) Erde, J.; Loo, R. R. O.; Loo, J. A. Enhanced FASP (eFASP) to Increase Proteome Coverage and Sample Recovery for Quantitative Proteomic Experiments. J. Proteome Res. 2014, 13 (4), 1885–1895. 10.1021/pr4010019.

(44) Rappsilber, J.; Mann, M.; Ishihama, Y. Protocol for Micro-Purification, Enrichment, Pre-Fractionation and Storage of Peptides for Proteomics Using StageTips. Nat. Protoc. 2007, 2 (8), 1896–1906. 10.1038/nprot.2007.261.

(45) Bacterial Vaccines: Methods and Protocols; Bidmos, F., Bossé, J., Langford, P., Eds.; Methods in Molecular Biology; Springer US: New York, NY, 2022; Vol. 2414. 10.1007/978-1-0716-1900-1.

(46) Wiśniewski, J. R.; Ostasiewicz, P.; Duś, K.; Zielińska, D. F.; Gnad, F.; Mann, M. Extensive Quantitative Remodeling of the Proteome between Normal Colon Tissue and Adenocarcinoma. Mol. Syst. Biol. 2012, 8 (1), 611. 10.1038/msb.2012.44.

(47) Shin, J.-B.; Krey, J. F.; Hassan, A.; Metlagel, Z.; Tauscher, A. N.; Pagana, J. M.; Sherman, N. E.; Jeffery, E. D.; Spinelli, K. J.; Zhao, H.; Wilmarth, P. A.; Choi, D.; David, L. L.; Auer, M.; Barr-Gillespie, P. G. Molecular Architecture of the Chick Vestibular Hair Bundle. Nat. Neurosci. 2013, 16 (3), 365–374. 10.1038/nn.3312.

(48) Tyanova, S.; Temu, T.; Cox, J. The MaxQuant Computational Platform for Mass Spectrometry-Based Shotgun Proteomics. Nat. Protoc. 2016, 11 (12), 2301–2319. 10.1038/nprot.2016.136.

(49) Teufel, F.; Almagro Armenteros, J. J.; Johansen, A. R.; Gíslason, M. H.; Pihl, S. I.; Tsirigos, K. D.; Winther, O.; Brunak, S.; Von Heijne, G.; Nielsen, H. SignalP 6.0 Predicts All Five Types of Signal Peptides Using Protein Language Models. Nat. Biotechnol. 2022, 40 (7), 1023–1025. 10.1038/s41587-021-01156-3.

(50) Kall, L.; Krogh, A.; Sonnhammer, E. L. L. Advantages of Combined Transmembrane Topology and Signal Peptide Prediction--the Phobius Web Server. Nucleic Acids Res. 2007, 35 (Web Server), W429–W432. 10.1093/nar/gkm256.

(51) Hallgren, J.; Tsirigos, K. D.; Pedersen, M. D.; Almagro Armenteros, J. J.; Marcatili, P.; Nielsen, H.; Krogh, A.; Winther, O. DeepTMHMM Predicts Alpha and Beta Transmembrane Proteins Using Deep Neural Networks; preprint; Bioinformatics, 2022. 10.1101/2022.04.08.487609.

(52) Fimereli, D. K.; Tsirigos, K. D.; Litou, Z. I.; Liakopoulos, T. D.; Bagos, P. G.; Hamodrakas, S. J. CW-PRED: A HMM-Based Method for the Classification of Cell Wall-Anchored Proteins of Gram-Positive Bacteria. In Artificial Intelligence: Theories and Applications; Maglogiannis, I., Plagianakos, V., Vlahavas, I., Eds.; Hutchison, D., Kanade, T., Kittler, J., Kleinberg, J. M., Mattern, F., Mitchell, J. C., Naor, M., Nierstrasz, O., Pandu Rangan, C., Steffen, B., Sudan, M., Terzopoulos, D., Tygar, D., Vardi, M. Y., Weikum, G., Series Eds.; Lecture Notes in Computer Science; Springer Berlin Heidelberg: Berlin, Heidelberg, 2012; Vol. 7297, pp 285–290. 10.1007/978-3-642-30448-4_36.

(53) Hernández-Plaza, A.; Szklarczyk, D.; Botas, J.; Cantalapiedra, C. P.; Giner-Lamia, J.; Mende, D. R.; Kirsch, R.; Rattei, T.; Letunic, I.; Jensen, L. J.; Bork, P.; von Mering, C.; Huerta-Cepas, J. eggNOG 6.0: Enabling Comparative Genomics across 12 535 Organisms. Nucleic Acids Res. 2023, 51 (D1), D389–D394. 10.1093/nar/gkac1022.

(54) Mistry, J.; Chuguransky, S.; Williams, L.; Qureshi, M.; Salazar, G. A.; Sonnhammer, E. L. L.; Tosatto, S. C. E.; Paladin, L.; Raj, S.; Richardson, L. J.; Finn, R. D.; Bateman, A. Pfam: The Protein Families Database in 2021. Nucleic Acids Res. 2021, 49 (D1), D412–D419. 10.1093/nar/gkaa913.

(55) Paysan-Lafosse, T.; Blum, M.; Chuguransky, S.; Grego, T.; Pinto, B. L.; Salazar, G. A.; Bileschi, M. L.; Bork, P.; Bridge, A.; Colwell, L.; Gough, J.; Haft, D. H.; Letunić, I.; Marchler-Bauer, A.; Mi, H.; Natale, D. A.; Orengo, C. A.; Pandurangan, A. P.; Rivoire, C.; Sigrist, C. J. A.; Sillitoe, I.; Thanki, N.; Thomas, P. D.; Tosatto, S. C. E.; Wu, C. H.; Bateman, A. InterPro in 2022. Nucleic Acids Res. 2023, 51 (D1), D418–D427. 10.1093/nar/gkac993.

(56) Szklarczyk, D.; Franceschini, A.; Wyder, S.; Forslund, K.; Heller, D.; Huerta-Cepas, J.; Simonovic, M.; Roth, A.; Santos, A.; Tsafou, K. P.; Kuhn, M.; Bork, P.; Jensen, L. J.; von Mering, C. STRING V10: Protein–Protein Interaction Networks, Integrated over the Tree of Life. Nucleic Acids Res. 2015, 43 (D1), D447–D452. 10.1093/nar/gku1003.

(57) Kim, B. J.; Park, J. H.; Park, T. H.; Bronstein, P. A.; Schneider, D. J.; Cartinhour, S. W.; Shuler, M. L. Effect of Iron Concentration on the Growth Rate of *Pseudomonas Syringae* and the Expression of Virulence Factors in *Hrp*-Inducing Minimal Medium. Appl. Environ. Microbiol. 2009, 75 (9), 2720–2726. 10.1128/AEM.02738-08.

(58) Cerdeno-Tarraga, A. M. The Complete Genome Sequence and Analysis of Corynebacterium Diphtheriae NCTC13129. Nucleic Acids Res. 2003, 31 (22), 6516–6523. 10.1093/nar/gkg874.

(59) Greenfield, L.; Bjorn, M. J.; Horn, G.; Fong, D.; Buck, G. A.; Collier, R. J.; Kaplan, D. A. Nucleotide Sequence of the Structural Gene for Diphtheria Toxin Carried by Corynebacteriophage Beta. Proc. Natl. Acad. Sci. 1983, 80 (22), 6853–6857. 10.1073/pnas.80.22.6853.

(60) Cox, J.; Mann, M. MaxQuant Enables High Peptide Identification Rates, Individualized p.p.b.-Range Mass Accuracies and Proteome-Wide Protein Quantification. Nat. Biotechnol. 2008, 26 (12), 1367–1372. 10.1038/nbt.1511.

(61) Dreisbach, A.; Wang, M.; van der Kooi-Pol, M. M.; Reilman, E.; Koedijk, D. G. A. M.; Mars, R. A. T.; Duipmans, J.; Jonkman, M.; Benschop, J. J.; Bonarius, H. P. J.; Groen, H.; Hecker, M.; Otto, A.; Bäsell, K.; Bernhardt, J.; Back, J. W.; Becher, D.; Buist, G.; van Dijl, J. M. Tryptic Shaving of *Staphylococcus Aureus* Unveils Immunodominant Epitopes on the Bacterial Cell Surface. J. Proteome Res. 2020, acs.jproteome.0c00043. 10.1021/acs.jproteome.0c00043.

(62) Olaya-Abril, A.; Gómez-Gascón, L.; Jiménez-Munguía, I.; Obando, I.; Rodríguez-Ortega, M. J. Another Turn of the Screw in Shaving Gram-Positive Bacteria: Optimization of Proteomics Surface Protein Identification in Streptococcus Pneumoniae. J. Proteomics 2012, 75 (12), 3733–3746. 10.1016/j.jprot.2012.04.037.

(63) Blum, M.; Chang, H.-Y.; Chuguransky, S.; Grego, T.; Kandasaamy, S.; Mitchell, A.; Nuka, G.; Paysan-Lafosse, T.; Qureshi, M.; Raj, S.; Richardson, L.; Salazar, G. A.; Williams, L.; Bork, P.; Bridge, A.; Gough, J.; Haft, D. H.; Letunic, I.; Marchler-Bauer, A.; Mi, H.; Natale, D. A.; Necci, M.; Orengo, C. A.; Pandurangan, A. P.; Rivoire, C.; Sigrist, C. J. A.; Sillitoe, I.; Thanki, N.; Thomas, P. D.; Tosatto, S. C. E.; Wu, C. H.; Bateman, A.; Finn, R. D. The InterPro Protein Families and Domains Database: 20 Years On. Nucleic Acids Res. 2021, 49 (D1), D344–D354. 10.1093/nar/gkaa977.

(64) Cox, J.; Hein, M. Y.; Luber, C. A.; Paron, I.; Nagaraj, N.; Mann, M. Accurate Proteome-Wide Label-Free Quantification by Delayed Normalization and Maximal Peptide Ratio Extraction, Termed MaxLFQ. Mol. Cell. Proteomics 2014, 13 (9), 2513–2526. 10.1074/mcp.M113.031591.

(65) Tyanova, S.; Temu, T.; Sinitcyn, P.; Carlson, A.; Hein, M. Y.; Geiger, T.; Mann, M.; Cox, J. The Perseus Computational Platform for Comprehensive Analysis of (Prote)Omics Data. Nat. Methods 2016, 13 (9), 731–740. 10.1038/nmeth.3901.

(66) Sharma, N. C.; Efstratiou, A.; Mokrousov, I.; Mutreja, A.; Das, B.; Ramamurthy, T. Diphtheria. Nat. Rev. Dis. Primer 2019, 5 (1), 81. 10.1038/s41572-019-0131-y.

(67) Liu, B.; Zheng, D.; Zhou, S.; Chen, L.; Yang, J. VFDB 2022: A General Classification Scheme for Bacterial Virulence Factors. Nucleic Acids Res. 2022, 50 (D1), D912–D917. 10.1093/nar/gkab1107.

(68) Peng, E. D.; Schmitt, M. P. Identification of Zinc and Zur-Regulated Genes in Corynebacterium Diphtheriae. PLOS ONE 2019, 14 (8), e0221711. 10.1371/journal.pone.0221711.

(69) Simpson-Lourêdo, L.; Silva, C. M. F.; Hacker, E.; Souza, N. F.; Santana, M. M.; Antunes, C. A.; Nagao, P. E.; Hirata, R.; Burkovski, A.; Villas Bôas, M. H. S.; Mattos-Guaraldi, A. L. Detection and Virulence Potential of a Phospholipase D-Negative Corynebacterium Ulcerans from a Concurrent Diphtheria and Infectious Mononucleosis Case. Antonie Van Leeuwenhoek 2019, 112 (7), 1055–1065. 10.1007/s10482-019-01240-4.

(70) Śmiga, M.; Ślęzak, P.; Wagner, M.; Olczak, T. Interplay between Porphyromonas Gingivalis Hemophore-Like Protein HmuY and Kgp/RgpA Gingipains Plays a Superior Role in Heme Supply. Microbiol. Spectr. 2023, 11 (2), e04593–22. 10.1128/spectrum.04593-22.

(71) Hou, M.; Huang, J.; Jia, T.; Guan, Y.; Yang, F.; Zhou, H.; Huang, P.; Wang, J.; Yang, L.; Dai, L. Deep Profiling of the Proteome Dynamics of Pseudomonas Aeruginosa Reference Strain PAO1 under Different Growth Conditions. J Proteome Res 2023.

(72) Azevedo, V.; D’Afonseca, V.; Ali; Santos, A.; Pinto; Magalhães; Faria; Barbosa; Guimarães; Eslabão; Almeida; Abreu; Neto; Carneiro; Cerdeira, L.; Ramos, R.; Hirata-Jr; Mattos-Guaraldi, A.; Trost; Tauch; Silva; Schneider; Miyoshi; Azevedo, V. Reannotation of the Corynebacterium Diphtheriae NCTC13129 Genome as a New Approach to Studying Gene Targets Connected to Virulence and Pathogenicity in Diphtheria. Open Access Bioinforma. 2012, 1. 10.2147/OAB.S25500.

(73) Griffin, M. E.; Klupt, S.; Espinosa, J.; Hang, H. C. Peptidoglycan NlpC/P60 Peptidases in Bacterial Physiology and Host Interactions. Cell Chem. Biol. 2023, 30 (5), 436–456. 10.1016/j.chembiol.2022.11.001.

(74) Gaday, Q.; Megrian, D.; Carloni, G.; Martinez, M.; Sokolova, B.; Ben Assaya, M.; Legrand, P.; Brûlé, S.; Haouz, A.; Wehenkel, A. M.; Alzari, P. M. FtsEX-Independent Control of RipA-Mediated Cell Separation in *Corynebacteriales*. Proc. Natl. Acad. Sci. 2022, 119 (50), e2214599119. 10.1073/pnas.2214599119.

(75) Peixoto, R. S.; Antunes, C. A.; Lourêdo, L. S.; Viana, V. G.; Santos, C. S. D.; Fuentes Ribeiro Da Silva, J.; Hirata Jr., R.; Hacker, E.; Mattos-Guaraldi, A. L.; Burkovski, A. Functional Characterization of the Collagen-Binding Protein DIP2093 and Its Influence on Host– Pathogen Interaction and Arthritogenic Potential of Corynebacterium Diphtheriae. Microbiology 2017, 163 (5), 692–701. 10.1099/mic.0.000467.

(76) Jacobitz, A. W.; Kattke, M. D.; Wereszczynski, J.; Clubb, R. T. Sortase Transpeptidases: Structural Biology and Catalytic Mechanism. Adv. Protein Chem. Struct. Biol. 2017, 109. 10.1016/bs.apcsb.2017.04.008.

(77) Dulberger, C. L.; Rubin, E. J.; Boutte, C. C. The Mycobacterial Cell Envelope — a Moving Target. Nat. Rev. Microbiol. 2020, 18 (1), 47–59. 10.1038/s41579-019-0273-7.

(78) Dautin, N.; de Sousa-d’Auria, C.; Constantinesco-Becker, F.; Labarre, C.; Oberto, J.; Li de la Sierra-Gallay, I.; Dietrich, C.; Issa, H.; Houssin, C.; Bayan, N. Mycoloyltransferases: A Large and Major Family of Enzymes Shaping the Cell Envelope of Corynebacteriales. Biochim. Biophys. Acta BBA - Gen. Subj. 2017, 1861 (1), 3581–3592. 10.1016/j.bbagen.2016.06.020.

(79) Kawashima, K.; Nagakubo, T.; Nomura, N.; Toyofuku, M. Iron Delivery through Membrane Vesicles in Corynebacterium Glutamicum. Microbiol. Spectr. 2023, 11 (3), e01222–23. 10.1128/spectrum.01222-23.

(80) Bou Raad, R.; Méniche, X.; De Sousa-d’Auria, C.; Chami, M.; Salmeron, C.; Tropis, M.; Labarre, C.; Daffé, M.; Houssin, C.; Bayan, N. A Deficiency in Arabinogalactan Biosynthesis Affects *Corynebacterium Glutamicum* Mycolate Outer Membrane Stability. J. Bacteriol. 2010, 192 (11), 2691–2700. 10.1128/JB.00009-10.

(81) Carel, C.; Marcoux, J.; Réat, V.; Parra, J.; Latgé, G.; Laval, F.; Demange, P.; Burlet-Schiltz, O.; Milon, A.; Daffé, M.; Tropis, M. G.; Renault, M. A. M. Identification of Specific Posttranslational *O*-Mycoloylations Mediating Protein Targeting to the Mycomembrane. Proc. Natl. Acad. Sci. 2017, 114 (16), 4231–4236. 10.1073/pnas.1617888114.

(82) Adindla, S.; Inampudi, K. K.; Guruprasad, K.; Guruprasad, L. Identification and Analysis of Novel Tandem Repeats in the Cell Surface Proteins of Archaeal and Bacterial Genomes Using Computational Tools. Comp. Funct. Genomics 2004, 5 (1), 2–16. 10.1002/cfg.358.

(83) Wästfelt, M.; Stålhammar-Carlemalm, M.; Delisse, A.-M.; Cabezon, T.; Lindahl, G. Identification of a Family of Streptococcal Surface Proteins with Extremely Repetitive Structure. J. Biol. Chem. 1996, 271 (31), 18892–18897. 10.1074/jbc.271.31.18892.

(84) Czibener, C.; Merwaiss, F.; Guaimas, F.; Del Giudice, M. G.; Serantes, D. A. R.; Spera, J. M.; Ugalde, J. E. BigA Is a Novel Adhesin of *Brucella* That Mediates Adhesion to Epithelial Cells: Adhesion of Brucella *to Epithelial Cells*. Cell. Microbiol. 2016, 18 (4), 500–513. 10.1111/cmi.12526.

(85) Draganova, E. B.; Akbas, N.; Adrian, S. A.; Lukat-Rodgers, G. S.; Collins, D. P.; Dawson, J. H.; Allen, C. E.; Schmitt, M. P.; Rodgers, K. R.; Dixon, D. W. Heme Binding by *Corynebacterium Diphtheriae* HmuT: Function and Heme Environment. Biochemistry 2015, 54 (43), 6598–6609. 10.1021/acs.biochem.5b00666.

(86) Draganova, E. B.; Adrian, S. A.; Lukat-Rodgers, G. S.; Keutcha, C. S.; Schmitt, M. P.; Rodgers, K. R.; Dixon, D. W. Corynebacterium Diphtheriae HmuT: Dissecting the Roles of Conserved Residues in Heme Pocket Stabilization. JBIC J. Biol. Inorg. Chem. 2016, 21 (7), 875–886. 10.1007/s00775-016-1386-3.

(87) Muraki, N.; Aono, S. Structural Basis for Heme Recognition by HmuT Responsible for Heme Transport to the Heme Transporter in *Corynebacterium Glutamicum*. Chem. Lett. 2016, 45 (1), 24–26. 10.1246/cl.150894.

(88) Wilks, A.; Schmitt, M. P. Expression and Characterization of a Heme Oxygenase (Hmu O) fromCorynebacterium Diphtheriae. J. Biol. Chem. 1998, 273 (2), 837–841. 10.1074/jbc.273.2.837.

(89) Kehl-Fie, T. E.; Skaar, E. P. Nutritional Immunity beyond Iron: A Role for Manganese and Zinc. Curr. Opin. Chem. Biol. 2010, 14 (2), 218–224. 10.1016/j.cbpa.2009.11.008.

(90) Peng, E. D.; Oram, D. M.; Battistel, M. D.; Lyman, L. R.; Freedberg, D. I.; Schmitt, M. P. Iron and Zinc Regulate Expression of a Putative ABC Metal Transporter in Corynebacterium Diphtheriae. J. Bacteriol. 2018, 200 (10). 10.1128/JB.00051-18.

(91) Bibb, L. A.; Schmitt, M. P. The ABC Transporter HrtAB Confers Resistance to Hemin Toxicity and Is Regulated in a Hemin-Dependent Manner by the ChrAS Two-Component System in Corynebacterium Diphtheriae. J. Bacteriol. 2010, 192 (18), 4606–4617. 10.1128/JB.00525-10.

(92) Bibb, L. A.; Kunkle, C. A.; Schmitt, M. P. The ChrA-ChrS and HrrA-HrrS Signal Transduction Systems Are Required for Activation of the *hmuO* Promoter and Repression of the *hemA* Promoter in *Corynebacterium Diphtheriae*. Infect. Immun. 2007, 75 (5), 2421–2431. 10.1128/IAI.01821-06.

(93) Schneitz, C.; Nuotio, L.; Lounatma, K. Adhesion of Lactobacillus Acidophilus to Avian Intestinal Epithelial Cells Mediated by the Crystalline Bacterial Cell Surface Layer (S-layer). J. Appl. Bacteriol. 1993, 74 (3), 290–294. 10.1111/j.1365-2672.1993.tb03028.x.

(94) Bahl, H.; Scholz, H.; Bayan, N.; Chami, M.; Leblon, G.; Gulik-Krzywicki, T.; Shechter, E.; Fouet, A.; Mesnage, S.; Tosi-Couture, E.; Gounon, P.; Mock, M.; Conway De Macario, E.; Macario, A. J. L.; Fernández-Herrero, L. A.; Olabarría, G.; Berenguer, J.; Blaser, M. J.; Kuen, B.; Lubitz, W.; Sára, M.; Pouwels, P. H.; Kolen, C. P. A. M.; Boot, H. J.; Palva, A.; Truppe, M.; Howorka, S.; Schroll, G.; Lechleitner, S.; Resch, S. IV. Molecular Biology of S-Layers. FEMS Microbiol. Rev. 1997, 20 (1–2), 47–98. 10.1111/j.1574-6976.1997.tb00304.x.

(95) Yeats, C.; Rawlings, N. D.; Bateman, A. The PepSY Domain: A Regulator of Peptidase Activity in the Microbial Environment? Trends Biochem. Sci. 2004, 29 (4), 169–172. 10.1016/j.tibs.2004.02.004.

(96) Rivera, M. Bacterioferritin: Structure, Dynamics, and Protein–Protein Interactions at Play in Iron Storage and Mobilization. Acc. Chem. Res. 2017, 50 (2), 331–340. 10.1021/acs.accounts.6b00514.

