## Supplementary material for "The exoproteome and surfaceome of toxigenic *Corynebacterium diphtheriae* 1737 and its response to iron-restriction and growth on human hemoglobin": Fig. S

[*]To whom correspondence should be addressed:

- **Figures S1-S6**

The following files are available free of charge.

- **Supplementary_Table_S1.xlsx**: The top 20 most abundant proteins from the cytoplasmic, supernatant, and envelope fractions and known *Corynebacterium* virulence factors.
- **Supplementary_Table_S2.xlsx**: Protein tables and quantification results for the exoproteome of *C. diphtheriae* strain *1737* under low-iron conditions.
- **Supplementary_Table_S3.xlsx**: Protein tables with quantification results for the surfaceome of *C. diphtheriae* strain *1737* under low-iron conditions.
- **Supplementary_Table_S4.xlsx**: Extracellular proteins that have a statistically significant change in secretion or surface-exposure in response to iron concentration or iron source.

**Table of Contents**


| Figure | Description | Page |
|---|---|---|
| **S1** | Evaluation of iron supplementation on *C. diphtheriae* growth | **3** |
| **S2** | Protein concentration quantification from culture fractions under low-iron conditions | **5** |
| **S3** | Localization profiles for the top 20 most abundant proteins from the supernatant, envelope, and cytoplasmic fractions under low-iron conditions | **6** |
| **S4** | Optimization of intact cell surface proteolysis under low-iron conditions | **7** |
| **S5** | Bioinformatic sequence motifs observed from proteins based on their assigned preferential secretion and surface-exposure levels under low-iron conditions. | **8** |
| **S6** | Localization profiles and abundance of putative heme-acquisition machinery from *C. diphtheriae* low-iron cultures | **10** |

**SUPPLEMENTARY FIGURES**

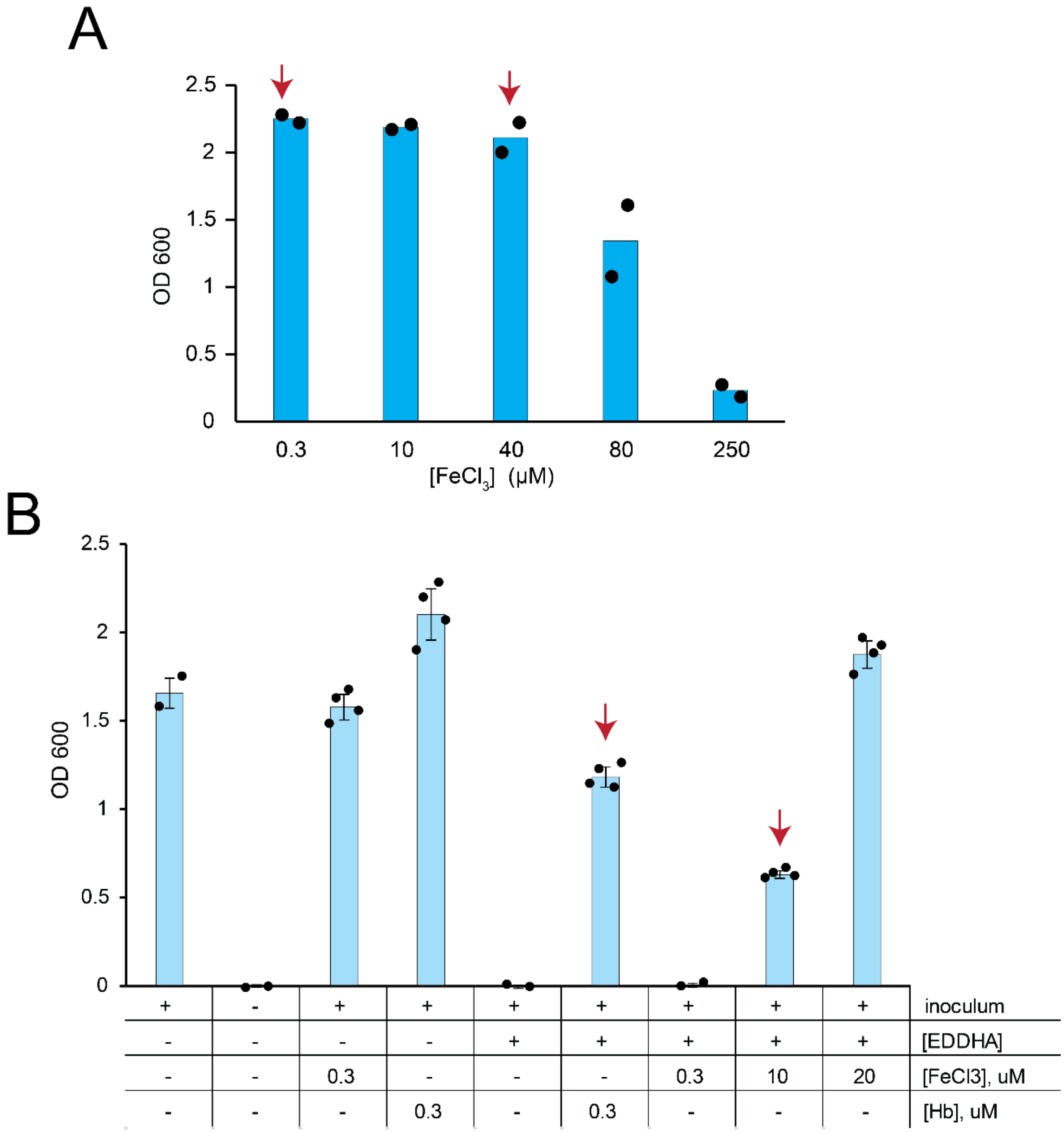


**Figure S1**: Preliminary growth studies of *C. diphtheriae* in iron-free mPGT media containing various supplementations to evaluate the effect of iron on cell growth. (A) Evaluation of various free iron ($FeCl_3$) supplementation concentrations in mPGT media effect on $OD_{600}$ from stationary phase growth. (B) Evaluating media supplements for optimization of iron-restricted culture conditions, with $OD_{600}$ values measured for cells at the late-exponential phase of their growth. Overnight cultures of *C. diphtheriae* grown in HIBTW (heart infusion broth supplemented with 0.2% tween-80) were diluted with two-parts (by volume) fresh HIBTW, cultured for 1-2 hours, washed with iron-free mPGT media, and resuspended in iron-free mPGT media. The resuspended cells were then used to as the inoculum to start cultures at an $OD_{600}$ of 0.1. The iron-chelator, EDDHA (ethylenediamine-N,N′-bis(2-hydroxyphenylacetic acid)), was supplemented at 10 μM in the indicated cultures. $FeCl_3$ and Hb were added to the cultures at differing concentrations as indicated. Red arrows indicate conditions 1 (A, left), 2a (A, right), 2b (B, left) and 2c (B, right) that were used in the proteomic experiments.

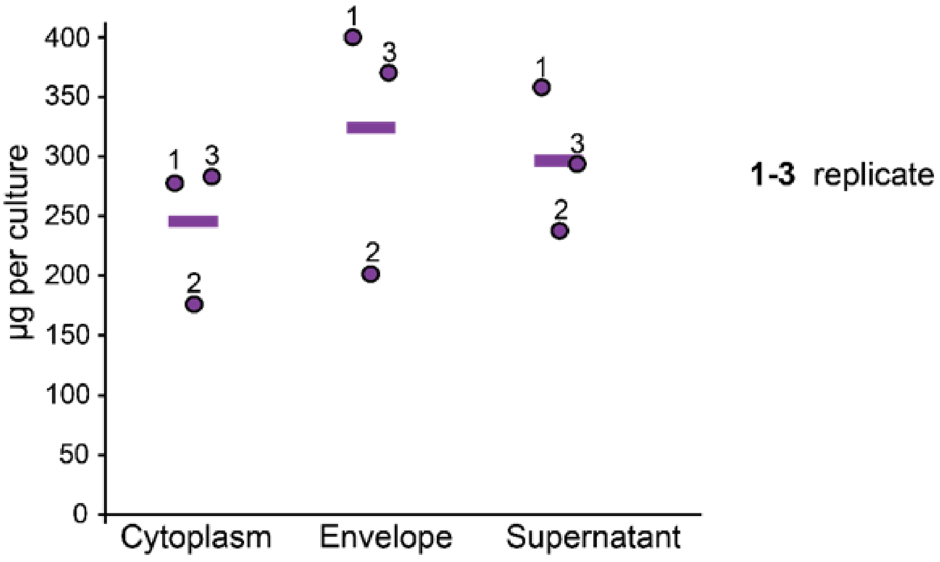


**Figure S2:** Absolute protein quantification results using the BCA assay for each culture fraction (supernatant, envelope, and cytoplasmic fraction) that was used to calculate the total culture protein content in *C. diphtheriae* 1737 (harvested at log-phase in low-iron conditions (condition 1)). Total protein quantification is described in the methods, and the results for each replicate are indicated by the numbers next to their respective datapoint. Protein concentrations in the cytoplasmic and cell envelope fractions were directly measured using the BCA assay from cell lysates. To measure protein concentrations in the secreted fraction culture supernatants were first dialyzed into PBS pH 7.4 (lysis buffer), carefully monitoring any volume changes, and then quantified using the BCA assay.

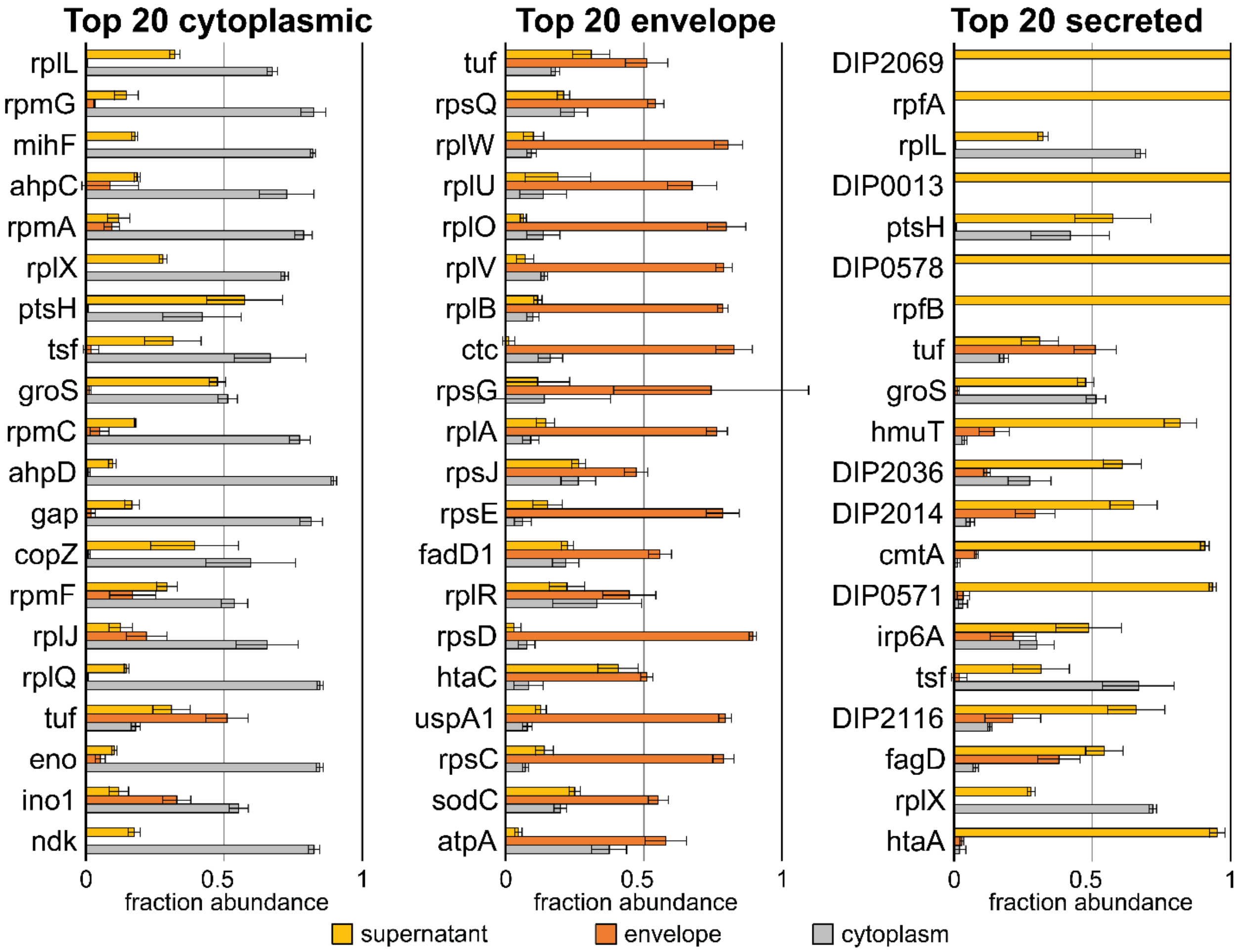


**Figure S3:** Localization profiles illustrating the relative subcellular partitioning for the top 20 most abundant proteins in the supernatant (left), envelope (middle), and cytoplasmic (right) fractions. Proteins are ordered by their relative abundance in each fraction, such that the most abundant protein is at the top. Data were acquired for cells grown in low iron media (condition 1). Standard deviations between the three biological replicates are shown for each protein.

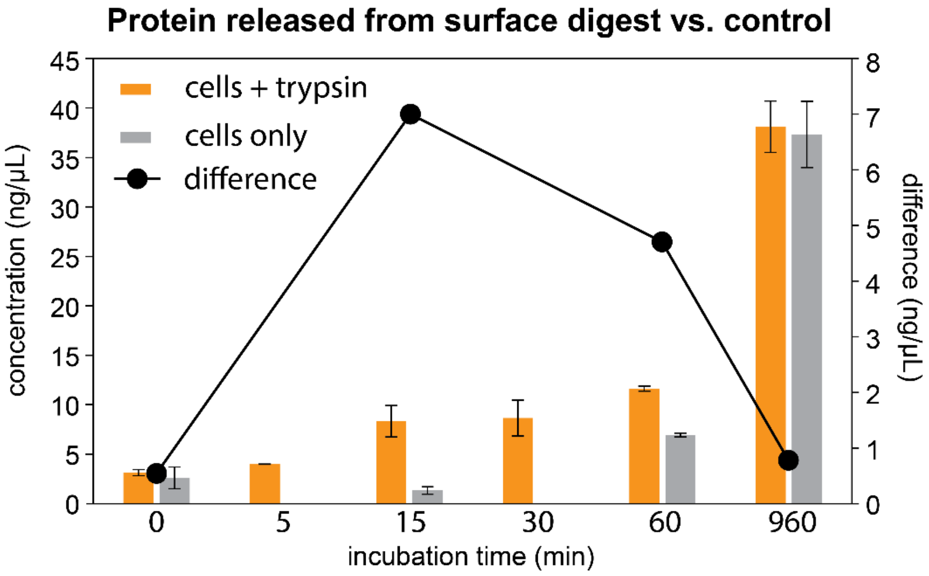


**Figure S4:** Optimization of trypsin digestion conditions to identify surface proteins in intact *C. diphtheriae* 1737 cells. Intact cells were incubated with either trypsin or a buffer control (50 mM trisHCl, pH 7.8) for varying amounts of time. The bar graph shows the concentration of released peptide determined by performing a BCA assay on the supernatant after the cells were proteolyzed for varying amounts of time. For each time point the difference between the released protein amount in the protease digested cells versus the no-protease control are represented by black dots (y-axis scale on the right indicates the difference). The largest difference occurred after 15 minutes. Therefore, this incubation time was employed in the protease digestion experiments.

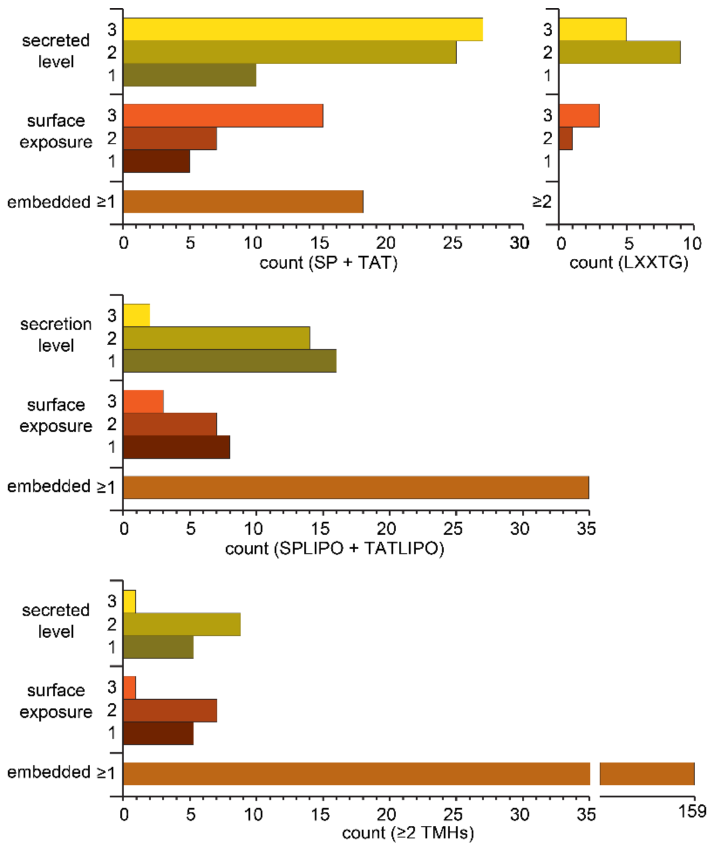


**Figure S5**: Bioinformatics analysis of secreted, surface-exposed, and envelope-embedded proteins identified by label-free quantitative MS analysis on log-phase low iron *C. diphtheriae* cultures. Plots showing the correspondence between the MS localization preferences and a bioinformatics analysis. The plot shows the number proteins whose location was determined by MS and their predicted location by bioinformatics. Secretion levels 1, 2, and 3 correspond to marginally, moderately, and predominantly secreted proteins (>30-50%, >50-90%, and >90% abundance from supernatant, respectively). Surface exposure categories of 1, 2, and 3 correspond to respectively marginally, moderately, and predominantly surface-exposed proteins proteins (>9.8%, >2.0-9.8% or >0.1-2.0% of their total envelope+cytoplasmic abundance released from a 15 minute proteolysis, respectively). For these proteins several bioinformatic programs were used to deduce their location. Proteins predicted to be secreted harbor Sec/SpI (Signal Peptidase

I) and TAT (twin arginine translocon) signal sequences, while extracellular membrane proteins contain Lipo/SpII (Signal peptidase II recognition and lipoylation sites) and TATLIPO (twin arginine motif transporter with SpII recognition) motifs. Proteins predicted to be covalently attached to the cell wall or embedded in the membrane contain “LXXTG” sorting signalsequences or > 2 transmembrane helices (> 2 TM), respectively. See methods for the programs used to predict protein location.

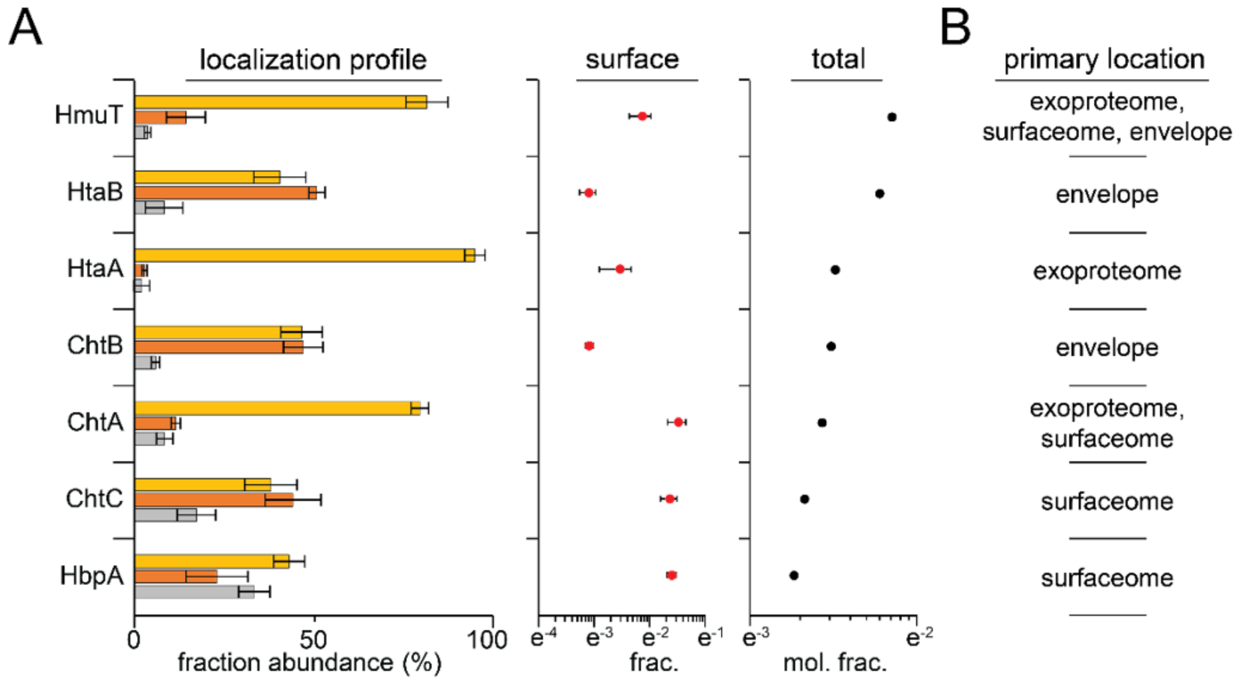


**Figure S6**: Localization of putative components of the *C. diphtheriae* heme-acquisition system under low-iron conditions. (A) Localization profiles (left), culture-normalized surface abundance (middle), and total culture abundance (supernatant, envelope, and cytoplasmic fractions) (right) for known heme-uptake components, ordered by total abundance (descending). Each bar in a protein's localization profile represents the percentage of that protein's total culture abundance that comes from the supernatant (yellow), envelope (orange), and cytoplasmic (grey) fractions. Relatively high cytoplasmic intensities from HbpA could arise from solubilization of the protein from the cell envelope during the lysis procedure (e.g. sheering from the cell surface), which is observed for highly abundant cell wall-anchored pilin and pilin-like surface proteins (**Supplementary Files S2**, **S3**). Culture-normalized surface digest abundances are in units of mass (calculated with intensities), while those of the total abundance are in molar units (calculated with iBAQ values[1]) (see methods). (B) Primary localization assignments are based on the protein's preferential localization (predominant > moderate > marginal) for secretion (“exoproteome”), surface-exposure (“surfaceome”), or envelope-embedding (“envelope”). ChtB, which was found with marginal preference across all fractions, was annotated with an envelope primary location because it is more abundant in the envelope relative to the supernatant and has both low surface abundance and surface-exposure (**Supplementary File S3**). Definitions of predominantly, moderately, and marginal preferential localization are defined in **Figure 2** and **Figure 3**.

### REFERENCES


(1) Cox, J.; Mann, M. MaxQuant Enables High Peptide Identification Rates, Individualized p.p.b.-Range Mass Accuracies and Proteome-Wide Protein Quantification. *Nat. Biotechnol.* **2008**, *26* (12), 1367–1372. https://doi.org/10.1038/nbt.1511.

(2) Teufel, F.; Almagro Armenteros, J. J.; Johansen, A. R.; Gíslason, M. H.; Pihl, S. I.; Tsirigos, K. D.; Winther, O.; Brunak, S.; Von Heijne, G.; Nielsen, H. SignalP 6.0 Predicts All Five Types of Signal Peptides Using Protein Language Models. *Nat. Biotechnol.* **2022**, *40* (7), 1023–1025. https://doi.org/10.1038/s41587-021-01156-3.

(3) Blum, M.; Chang, H.-Y.; Chuguransky, S.; Grego, T.; Kandasaamy, S.; Mitchell, A.; Nuka, G.; Paysan-Lafosse, T.; Qureshi, M.; Raj, S.; Richardson, L.; Salazar, G. A.; Williams, L.; Bork, P.; Bridge, A.; Gough, J.; Haft, D. H.; Letunic, I.; Marchler-Bauer, A.; Mi, H.; Natale, D. A.; Necci, M.; Orengo, C. A.; Pandurangan, A. P.; Rivoire, C.; Sigrist, C. J. A.; Sillitoe, I.; Thanki, N.; Thomas, P. D.; Tosatto, S. C. E.; Wu, C. H.; Bateman, A.; Finn, R. D. The InterPro Protein Families and Domains Database: 20 Years On. *Nucleic Acids Res.* **2021**, *49* (D1), D344–D354. https://doi.org/10.1093/nar/gkaa977.

(4) Cox, J.; Hein, M. Y.; Luber, C. A.; Paron, I.; Nagaraj, N.; Mann, M. Accurate Proteome-Wide Label-Free Quantification by Delayed Normalization and Maximal Peptide Ratio Extraction, Termed MaxLFQ. *Mol. Cell. Proteomics* **2014**, *13* (9), 2513–2526. https://doi.org/10.1074/mcp.M113.031591.

(5) Fimereli, D. K.; Tsirigos, K. D.; Litou, Z. I.; Liakopoulos, T. D.; Bagos, P. G.; Hamodrakas, S. J. CW-PRED: A HMM-Based Method for the Classification of Cell Wall-Anchored Proteins of Gram-Positive Bacteria. In *Artificial Intelligence: Theories and Applications*; Maglogiannis, I., Plagianakos, V., Vlahavas, I., Eds.; Hutchison, D., Kanade, T., Kittler, J., Kleinberg, J. M., Mattern, F., Mitchell, J. C., Naor, M., Nierstrasz, O., Pandu Rangan, C., Steffen, B., Sudan, M., Terzopoulos, D., Tygar, D., Vardi, M. Y., Weikum, G., Series Eds.; Lecture Notes in Computer Science; Springer Berlin Heidelberg: Berlin, Heidelberg, 2012; Vol. 7297, pp 285–290. https://doi.org/10.1007/978-3-642-30448-4_36.

(6) Hallgren, J.; Tsirigos, K. D.; Pedersen, M. D.; Almagro Armenteros, J. J.; Marcatili, P.; Nielsen, H.; Krogh, A.; Winther, O. *DeepTMHMM Predicts Alpha and Beta Transmembrane Proteins Using Deep Neural Networks*; preprint; Bioinformatics, 2022. https://doi.org/10.1101/2022.04.08.487609.

(7) Tyanova, S.; Temu, T.; Sinitcyn, P.; Carlson, A.; Hein, M. Y.; Geiger, T.; Mann, M.; Cox, J. The Perseus Computational Platform for Comprehensive Analysis of (Prote)Omics Data. *Nat. Methods* **2016**, *13* (9), 731–740. https://doi.org/10.1038/nmeth.3901.
